# *Plasmodium falciparum* quinine resistance is multifactorial and includes a role for the drug/metabolite transporters PfCRT and DMT1

**DOI:** 10.1101/2024.09.27.615529

**Authors:** Mariko Kanai, Sachel Mok, Tomas Yeo, Melanie J. Shears, Jin H. Jeon, Sunil K. Narwal, Leila S. Ross, Meseret T. Haile, Abhai K. Tripathi, Godfree Mlambo, Jonathan Kim, Eva Gil-Iturbe, John Okombo, Heekuk Park, Kharizta Wiradiputri, Kurt E. Ward, Felix D. Rozenberg, Kate J. Fairhurst, Talia S. Bloxham, Jessica L. Bridgford, Tanaya R. Sheth, Marcus C.S. Lee, Jennifer L. Small-Saunders, Filippo Mancia, Matthias Quick, Anne-Catrin Uhlemann, Photini Sinnis, David A. Fidock

**Author notes:** These authors contributed equally to this work (Tomas Yeo, Melanie J. Shears). Corresponding author’s.

## Abstract

The genetic basis of Plasmodium falciparum resistance to quinine (QN), a drug used to treat severe malaria, has long been enigmatic. To gain further insight, we used FRG-NOD human liver-chimeric mice to conduct a P. falciparum genetic cross between QN-resistant (Cam3.II) and QN-sensitive (NF54) parasites, which also differ in their susceptibility to chloroquine (CQ). By applying different selective conditions to progeny pools prior to cloning, we recovered 120 unique recombinant progeny. Drug profiling and quantitative trait loci analyses of the progeny revealed predominant peaks on chromosomes 7 and 12 associated with CQ and QN resistance, that is consistent with a multifactorial mechanism of resistance for these compounds. CQ and monodesethyl-CQ (md-CQ) resistance mapped to a chromosome 7 region harboring pfcrt as expected. However, for QN, resistance mapped to a dominant chromosome 7 peak centered 295 kb downstream of pfcrt, with pfcrt showing a smaller peak. We identified the drug/metabolite transporter 1 (DMT1) as the top chromosome 7 candidate due to its structural similarity to PfCRT and proximity to the peak. Deleting DMT1 in QN-resistant Cam3.II parasites significantly sensitized the parasite to QN but not to the other drugs tested, suggesting that DMT1 mediates QN response specifically. We localized DMT1 to structures associated with vesicular trafficking, as well as the parasitophorous vacuolar membrane, lipid bodies, and the digestive vacuole. We also observed that mutant DMT1 transports more QN than the wild-type isoform in vitro. Gene editing confirmed an additional role for mutant PfCRT in mediating QN resistance. In addition, we identified an ATP-dependent zinc metalloprotease (FtsH1) as one of the top candidates in the chromosome 12 locus and confirmed its role as a potential mediator of QN resistance and a modulator of md-CQ resistance using CRISPR/Cas9 SNP-edited lines. Interestingly, this chromosome 12 region mapped to resistance to both CQ and QN and was preferentially co-inherited with pfcrt. Our study demonstrates that DMT1 is a novel marker of QN resistance and that a new chromosome 12 locus associates with CQ and QN response, with ftsh1 as a potential candidate, suggesting these genes in addition to pfcrt should be genotyped in surveillance and clinical settings.

## Introduction

In 2023, there were an estimated 263 million cases of malaria, and an estimated 597,000 deaths, mostly African children^1^. Treatment of *Plasmodium falciparum* uncomplicated malaria depends on artemisinin-based combination therapies (ACTs), which combine fast-acting yet rapidly cleared artemisinin (ART) derivatives with longer-lasting partner drugs (mostly lumefantrine or amodiaquine). Artesunate is also the first-line treatment for severe, life-threatening malaria.

Alarmingly, *P. falciparum* parasites have now acquired partial resistance to ART derivatives in multiple African countries, having earlier saturated the greater Mekong Subregion in Southeast Asia^2,3^. Given the potential for widespread *P. falciparum* resistance to ARTs in Africa and the possible loss of ACT efficacy, alternative drugs with distinct modes of action are urgently needed. Trials are underway with several candidate drugs^4^, however their rollout is years away. A potential alternative is to reintroduce licensed drugs such as QN, including for treating severe malaria. QN is also recommended (alone or with clindamycin) to treat *P. falciparum* malaria infections in pregnant women during the first trimester^1,5^. This drug has largely retained clinical efficacy since its first use in the 1600s, with resistance being slow to develop. QN resistance was first reported in ∼1910 in Brazil, with subsequent reports from areas in South America, Africa, and Southeast Asia^6^. In these cases, QN resistance was usually low-grade, with QN having a delayed or reduced effect. QN treatment failure may result from parasite resistance^7^ or other factors including variability in QN pharmacokinetics, compliance, and drug quality.

Here, we sought to gain a better understanding of genetic factors associated with QN resistance. This mechanism is thought to be multifactorial and has been partially associated with sequence changes in the *P. falciparum* chloroquine resistance transporter (*pfcrt)* (via single nucleotide polymorphisms (SNPs)), multidrug resistance protein 1 (*pfmdr1)* (both copy number and SNPs), sodium/hydrogen exchanger (*pfnhe-1)* (DNNND repeat polymorphisms), and the HECT E3 ubiquitin ligase *pfut* (SNPs)^8–13^. *pfnhe-1* and *pfut*, however, are minor strain-specific determinants at best, suggesting the involvement of other genetic loci. QN, mefloquine (MFQ), and lumefantrine (LMF) are all classified as aryl-amino alcohols (**Extended Data 1a**) and appear to share partially overlapping modes of action, although the exact mechanisms remain unclear^14^.

We also explored CQ, a 4-aminoquinoline (**Extended Data 1a**) and former first-line drug for *P. falciparum* that is still used to treat *P. vivax* malaria. CQ acts on the heme detoxification pathway by diffusing into the acidic digestive vacuole (DV), where it is protonated and binds to heme. This binding prevents heme detoxification into hemozoin (Hz), with free heme and heme-CQ adducts killing the parasite. PfCRT is the primary determinant of CQ resistance, with a minor contribution from mutant PfMDR1, which, like PfCRT, is a DV membrane transporter^10,15–19^. Both mutant *pfcrt* and *pfmdr1* are selected against in favor of their wild-type alleles in regions that use artemether-lumefantrine, the major ACT across Africa^20–22^.

Genetic crosses are a powerful forward genetics tool to investigate the molecular basis of a variety of phenotypes including drug resistance. These studies have identified mutations in *dhps, dhfr* and *pfcrt* as primary determinants of resistance to sulfadoxine, pyrimethamine and CQ, respectively^23–26^, and have been applied to study ART resistance mechanisms^27,28^. This methodology has been empowered by the development of a human liver-chimeric mouse model that obviates the earlier requirement for splenectomized monkeys^29^. Here, we applied this model to explore *P. falciparum* resistance to QN and CQ.

## Results

### Implementing a genetic cross to map markers of QN and CQ resistance

To identify markers of QN resistance, we implemented a genetic cross between Cam3.II and NF54 parasites (**Fig.1a**). Cam3.II is resistant to QN and CQ and originates from Southeast Asia^30,31^. This parasite expresses genetic variants of *pfcrt* and *pfut* that have been partially associated with QN resistance, as well as the *pfmdr1* N86/Y184F variant that does not associate with QN^10,13,18,32^. Cam3.II also harbors two copies of *pfmdr1* (**Fig. 1a**). NF54 is sensitive to both drugs and is wild-type at these three single-copy genes. Mean QN and CQ IC_50_ values (in media with human serum) for Cam3.II were ∼90 nM, and for NF54 were ∼10 nM. Based on our whole-genome sequence analysis, Cam3.II and NF54 differ at 13,116 SNP positions, providing ample resolution to genetically map primary and secondary mediators of resistance to both drugs.

**Fig. 1.**
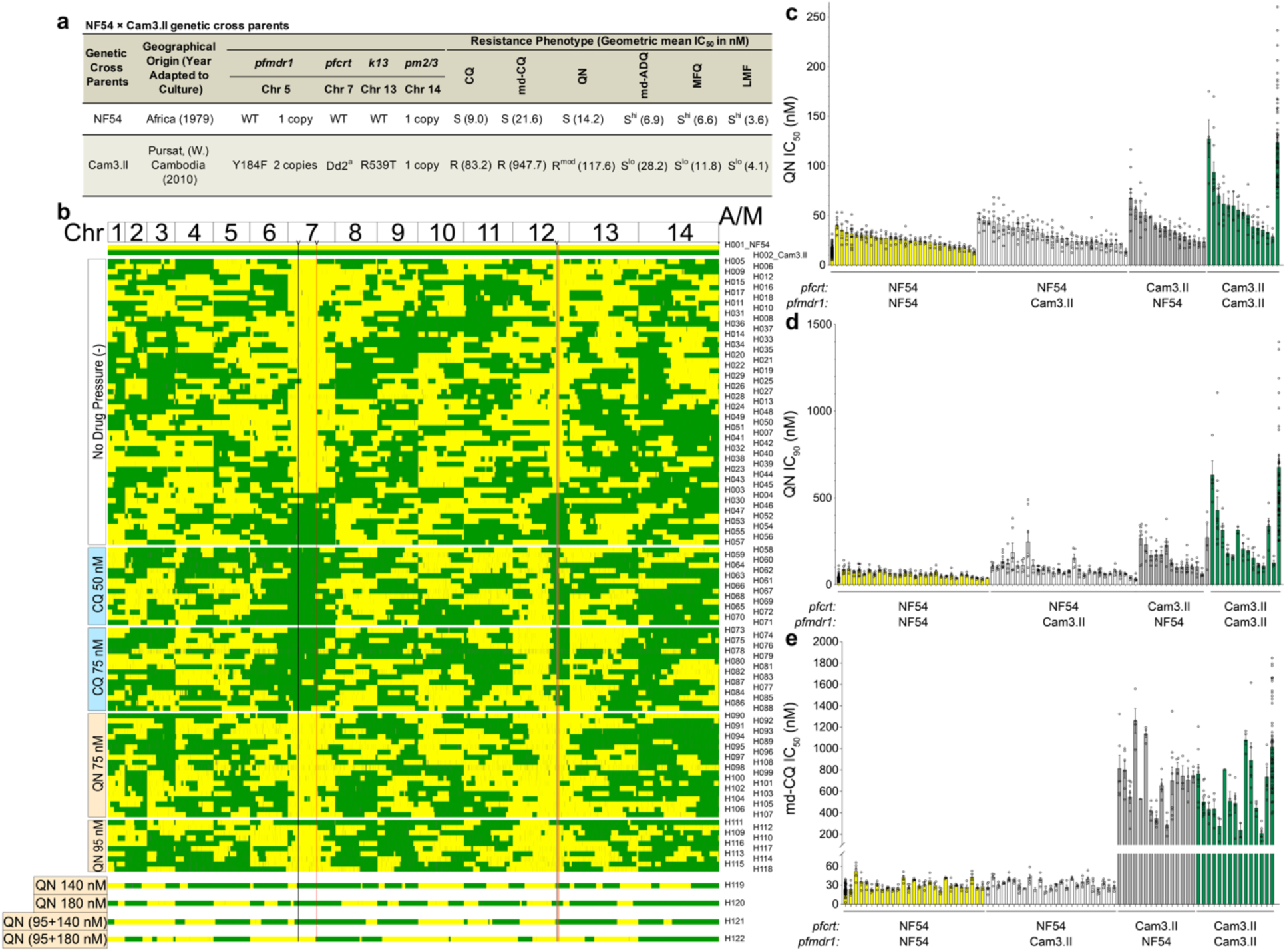
A *P. falciparum* NF54×Cam3.II genetic cross yielded 120 independent recombinant progeny with diverse genomic and drug resistance profiles. **a**, NF54×Cam3.II genetic cross parent information. ^a^Dd2 PfCRT haplotype: M74I/N75E/K76T/A220S/Q271E/N326S/I356T/R371I. *k13*: kelch13 (PF3D7_1343700); *pfcrt*: chloroquine resistance transporter (PF3D7_0709000); *pfmdr1*: multidrug resistance protein 1 (PF3D7_0523000). ADQ: amodiaquine; ART: artemisinin; CQ: chloroquine; LMF: lumefantrine; md-ADQ: mono-desethyl amodiaquine; md-CQ: monodesethyl-chloroquine; MFQ: mefloquine; md-CQ: monodesethyl-chloroquine; PPQ: piperaquine; QN: quinine. R, resistant; R^mod^, moderate resistance; S, sensitive; S^hi^, high sensitivity; S^lo^, low sensitivity. Geometric mean IC_50_ values in nM are listed in the parentheses (N=4−85 with technical duplicates). **b**, Recombinant progeny representing each of 120 unique haplotypes were hierarchically clustered by similarity based on their NF54 (yellow) or Cam3.II (green) alleles at 13,116 SNP positions within each selection condition: no drug pressure, chloroquine (CQ) 50 nM or 75 nM, or quinine (QN) 75 nM, 95 nM, 140 nM, 180 nM, (95nM+140nM), (95nM+180nM), or (95nM+140nM+240nM), in that order. Some haplotypes were obtained in more than one selection condition (**Table S4**). NF54 (H001) and Cam3.II (H002) parents are shown on top. H, haplotype; A/M, apicoplast/mitochondria (these were always coinherited); black line, *pfcrt*; red line, *dmt1;* grey line*, ftsh1*; red box, QN IC_90_, md-CQ IC_90_, CQ IC_50_ and IC_90_ chromosome 12 QTL locus. **c-e**, Mean ± SEM QN IC_50_ and IC_90_ values are shown in **c** and **d**, respectively, and md-CQ IC_50_ values are shown in **e** (N,n=1−85, with a majority of >4, with technical duplicates). Data were generated with parasites cultured in media with Albumax (no serum).

To achieve this cross, we mixed mature NF54 and Cam3.II gametocytes at a 1:1 ratio and fed them to female *Anopheles stephensi* mosquitoes^33^ (**Extended Data 2a**). In the midgut, gamete formation and subsequent sexual reproduction and recombination can generate thousands of haploid sporozoites that migrate to the salivary glands, ready to infect a mammalian host. Following parasitized blood feeds, we observed an infection prevalence of ∼85%, with ∼20 oocysts resulting in ∼40,000 salivary gland sporozoites per mosquito (**Table S1**). We then infected four FRG NOD human liver-chimeric (huHep) mice with ∼60,000 to 200,000 progeny sporozoites per mouse. Parasites were inoculated via mosquito bite or intravenously. After six days of liver-stage development, human red blood cells (RBCs) were infused and on day seven, blood was drawn to capture haploid asexual blood stage (ABS) parasites. These progeny were cultured *in vitro* for one to two 48 h generations of intra-erythrocytic development and replication before cryopreservation.

### Combining bulk progeny cloning with differential drug selections yields 120 unique recombinant progeny

We thawed progeny bulk freezes from the four mice (A−D), expanded these cultures in serum-containing media without drug pressure (‘no drug bulks’), or under various CQ and QN concentrations for 3–4 days, and cloned the progeny by limiting dilution (**Extended Data 2b,c**). Recombinants were identified by SNP and microsatellite (MS) genotyping and confirmed by whole-genome sequencing.

For each progeny, we classified alleles at each of the 13,116 SNP positions (**Table S2**) that distinguish the parents. A subset of 1,836 SNPs was then used to generate genetic maps, yielding a single linkage group for each of the 14 chromosomes. The recombination rate of 13.7kb/cM was consistent with prior crosses^28,29,34–37^ (**Table S3**). In total, the progeny segregated into 120 unique recombinant haplotypes (**Table S4** and **Fig. 1b**). 55 were obtained without drug pressure prior to cloning. Of these, 45 (82%) were wild-type for *pfcrt*, consistent with mutant *pfcrt* exerting a fitness cost in the absence of positive selection. Pressuring with various concentrations of CQ or QN pressure yielded an additional 65 haplotypes (**Fig. 1b**). Some QN selections included two to three rounds with increasing concentrations, yielding four of the 65 unique haplotypes, suggesting that high-grade selection can help to recover rare recombinants (and potentially those with a fitness cost). Only 5 unique recombinant haplotypes were represented by progeny obtained in both the CQ and QN selections, suggesting non-identical mechanisms of resistance to these drugs.

Most recombinant progeny did not show any major allele frequency skews across their genomes that deviated from the expected ∼40−60% parental allele frequencies (**Extended Data 3** and **Table S4**). As expected, all CQ-selected progeny possessed mutant *pfcrt* from the Cam3.II parent (matching the well-characterized Dd2 allele; **Supplemental Fig. 1a**). We obtained 26 haplotypes from progeny bulk pools selected with QN at 95 nM (∼1× QN IC_50_ for Cam3.II) or higher concentrations, of which 17 (65%) had wild-type (WT) *pfcrt* and 15 (58%) had WT *dmt1* (**Supplemental Fig. 1b**). We noted some enrichment for mutant *pfcrt* in parasites selected with QN (**Supplemental Fig. 1a**), suggesting a role for this chromosomal region in QN resistance.

### Bulk segregant analysis identifies two quantitative trait loci peaks on chromosome 7 that associate with quinine resistance

We then applied bulk segregant analysis (also known as linkage group selection analysis) to determine genetic loci associated with QN resistance. This approach employed QN pressure on bulk progeny pools, followed by whole-genome sequencing to measure allele frequency changes and identify genetic loci associated with the trait of interest. By pressuring four bulk cultures for 4 days with different concentrations of QN (∼1.5, 2.0, or 2.6× IC_50_ of the Cam3.II parent), with or without an initial period of exposure to QN at 1× IC_50_, we obtained 33 parasite pools. Those DNAs were then whole-genome sequenced. We also sequenced DNA from mock-treated bulk cultures processed in parallel (**Extended Data 2b,d** and **Table S5**).

We then compared the genome data of bulk progeny pools from QN-treated parasites vs ‘no drug timepoint control’ or day 0 (D0) control samples. Pairwise comparisons between two different QN selections and their ‘D0’ controls yielded largely consistent quantitative trait loci (QTLs) (**Extended Data 4**). The comparison using the higher QN concentration (180 nM) yielded a significant chromosome 7 peak, spanning 58 kb, that was located 191 kb downstream of *pfcrt* and that was driven by QN-resistant Cam3.II loci (**Table S6**). This 58 kb region did not contain any previously known drug resistance markers. These results show the power of bulk segregant analysis as a screening tool to identify regions of interest with multigenic traits. Bulk segregant analysis, however, needs to be followed-up by progeny clone-based QTL analysis to narrow down candidates and reveal epistatic relationships.

### Progeny clone-based trait mapping identifies novel resistance-associated peaks on chromosomes 7 and 12

To further resolve the segments associated with QN resistance, we implemented QTL analysis with individually cloned progeny (**Fig. 2e** and **Table S7**) by measuring their QN IC_50_ and IC_90_ values in 72-hour assays (**Fig. 1c,d** and **Extended Data 5a** and **Table S4**). These phenotypic data were then compared with their whole-genome sequences (**Fig. 2a,b,d,e** and **Extended Data 5b,c**). As a comparator, we also included CQ and md-CQ.

**Fig. 2.**
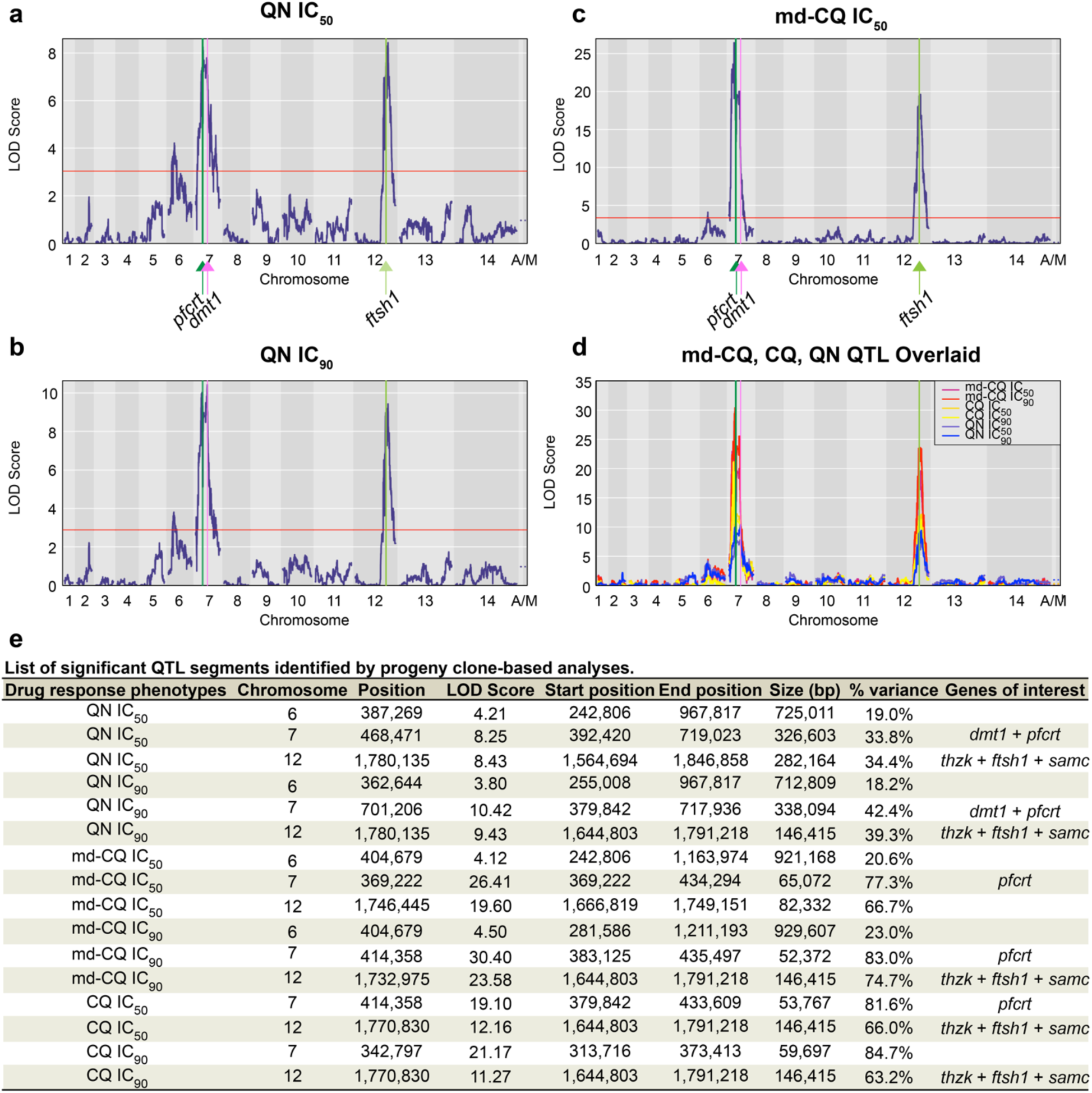
QTL mapping of QN, CQ, and md-CQ progeny reveals a non-*pfcrt*-centered chromosome 7 QN IC_90_ peak and an additional shared peak on chromosome 12. **a-c**, LOD plots for mean QN IC_50_ (**a**), QN IC_90_ (**b**), and md-CQ IC_50_ (**c**) values. The red line indicates the 95% probability threshold. **d**, md-CQ, CQ, and QN LOD plots overlaid. QTL, quantitative trait locus; LOD, log of the odds; md-CQ, monodesethyl-chloroquine; CQ, chloroquine; QN, quinine; IC_50_ and IC_90_, 50% and 90% growth inhibitory concentrations. **e**, List of significant QTL segments identified by progeny clone-based analysis. chromosome 7: *pfcrt* (dark green line), *dmt1* (pink line). chromosome 12: *thzk* and *ftsh1* (light green line).

For QN, we obtained the highest, statistically significant peaks on chromosomes 7 and 12, and several other minor QTLs with lower LOD scores, confirming that QN resistance is multigenic (**Fig. 2a,b,d**). The chromosome 12 QN QTLs either encompassed (IC_50_) or perfectly matched (IC_90_) the CQ and md-CQ QTLs. Strikingly, whereas the QN IC_50_ chromosome 7 peak centered close to *pfcrt*, the QN IC_90_ (high-grade resistance) peak was centered 296 kb downstream of *pfcrt* and encompassed 33 genes with nonsynonymous SNPs between the parents (**Fig. 2e** and **Table S7**).

For md-CQ and CQ, we found two shared major peaks—a chromosome 7 peak harboring *pfcrt* and a chromosome 12 peak that was usually co-inherited with *pfcrt* (i.e. mostly mutant or WT at both loci), with evidence of strong linkage disequilibrium (**Fig. 2c,e** and **Extended Data 5** and **Table S4**). The md-CQ IC_90_ and CQ IC_50_ and IC_90_ chromosome 12 peaks consist of 30 genes, of which 10 have nonsynonymous SNPs between the parents. These SNPs were also observed in field populations (**Table S7**). A smaller chromosome 6 peak observed with QN and to a marginal extent md-CQ was 245kb (QN IC_50_ and IC_90_), 49kb (md-CQ IC_50_), or 2kb (md-CQ IC_90_) away from the *aat1* S258L/F313S variant that was recently proposed to be an additional marker of CQ resistance^38^.

### The chromosome 12 peak harbors *samc*, *thzk*, and *ftsh1* that associate with quinine and chloroquine resistance

From amongst the 10 genes on chromosome 12 that harbored point mutations, we selected *samc* (PF3D7_1241600), *ftsh1* (PF3D7_1239700), and *thzk* (PF3D7_1239600) as top candidates (**Fig. 3**) based on their previous associations with *P. falciparum* resistance to antiplasmodial compounds^39–41^. In an earlier transposon mutagenesis screen with *in vitro* cultured *P. falciparum* ABS parasites^42^, *samc* was predicted to be essential, whereas *ftsh1* and *thzk* were predicted to be dispensable with either no or slight fitness cost, respectively.

**Fig. 3.**
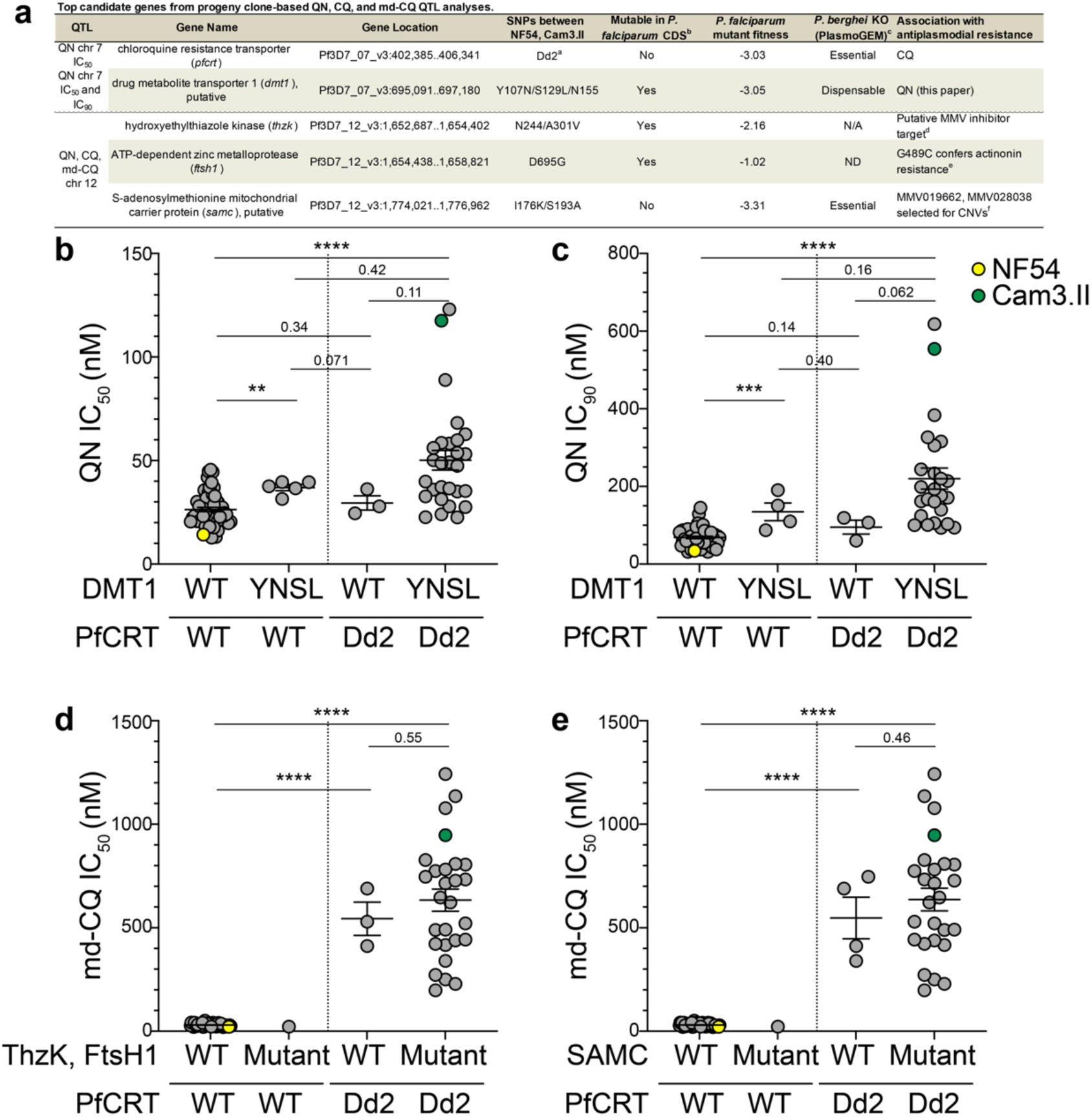
Identification of the top candidate genes from the QN chromosome 7 and QN, CQ, md-CQ chromosome 12 QTL peaks. **a**, Top candidate genes from progeny clone-based QN, CQ, and md-CQ QTL analyses. All genes are expressed in the *P. falciparum* ABS parasites^100^. QTL, quantitative trait locus; SNP, single-nucleotide polymorphism; QN, quinine; CQ, chloroquine; md, monodesethyl; CDS, coding sequence; KO, knockout; CNV, copy number variants; ND, not determined; N/A, not applicable. ^a^Dd2 PfCRT haplotype: M74I/N75E/K76T/A220S/Q271E/N326S/I356T/R371I. ^b^ref. 42, the closer the mutant fitness score is to -4, the higher the predicted fitness cost. ^c^ref. 101. ^d^Personal communication with Dr. Susan Wyllie. ^e^ref. 41. ^f^ref. 40. **b**-**e**, QN (**b**,**c**) and md-CQ. **d**,**e**, IC levels for the profiled unedited cross progeny and two parents, grouped by *pfcrt*, *dmt1*, *thzk*, *ftsh1*, and *samc* genotypes. Statistical significance was determined by Mann-Whitney *U* tests (N,n=1−27, with technical duplicates). \*\**p* < 0.01, \*\*\**p* < 0.001, \*\*\**p* < 0.0001.

We also looked for evidence of selective pressures on these genes in *P. falciparum* field isolates, using the Pf3k genome database^43^ (**Extended Data 6b-d**). FtsH1 D695G (present in Cam3.II) was observed in Southeast Asia and not in Africa. Haplotypes encoding the SAMC mutations I176K/R (K observed in Cam3.II) and S193A/P/T (A observed in Cam3.II) were prevalent, especially in Southeast Asia. The Cam3.II ThzK A301V mutation was present at low prevalence across Southeast Asian and African countries. One explanation for why these loci would not have been detected in prior CQ QTL analyses with the HB3×Dd2 and 7G8×GB4 genetic crosses is that the *ftsh1, thzk* and *samc* alleles were identical between those parents.

### FtsH1 mutations can modulate quinine and chloroquine resistance

To directly test whether the chromosome 12 QTL candidates *samc, thzk* or *ftsh1* contribute to QN and CQ resistance, we reverted these mutant alleles to WT in Cam3.II parasites (**Supplemental Fig. 2** and **Tables S8, S9**). We then tested for susceptibility to QN, md-CQ, and CQ in gene-edited lines (denoted as *samc*^WT^, *thzk*^WT^ or *ftsh1*^WT^) compared to the unedited Cam3.II parent (**Fig. 4a-f** and **Table S10**). *samc*^WT^ and *thzk*^WT^ parasites showed no significant difference to all three drugs when compared to Cam3.II. Also, no shift was observed when introducing the *thzk* A301V mutation into NF54 parasites (**Table S10**). In contrast, reverting *ftsh1* to WT in Cam3.II caused a significant decrease in the level of resistance to QN, CQ and md-CQ, with IC_50_ decreases averaging 33% to 50%. These results provide evidence that *ftsh1* may act as a potential contributor to QN resistance and mutant *pfcrt*-mediated CQ and md-CQ resistance.

**Fig. 4.**
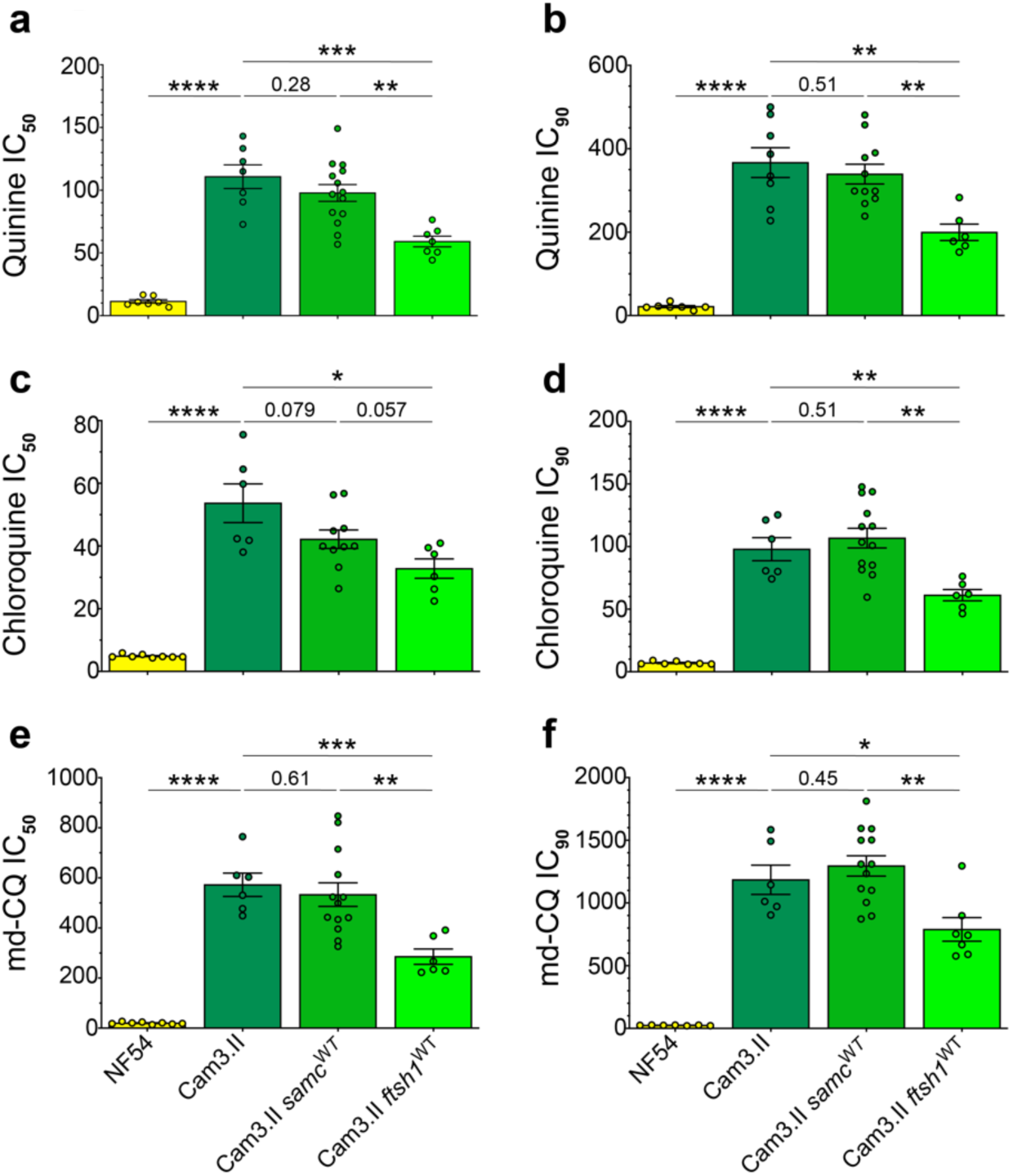
*ftsh1* is a candidate gene for the chromosome 12 QTL locus. **a**-**f**, Quinine (**a**,**b**), chloroquine (**c**,**d**), and md-Chloroquine (**e**,**f**) response in *samc* and *ftsh1* SNP-edited Cam3.II as measured by mean ± SEM IC_50_ or IC_90_ values. For each parasite, the parental unedited parasite (endogenous haplotype) (dark color) and the SNP-edited parasite (wild-type revertant parasite) (light color) were profiled. *p*-values (Student’s *t*-test) are indicated (N,n=6−16,2). \**p* < 0.05; \*\**p* < 0.01; \*\*\**p* < 0.001.

### *dmt1* is a top candidate for mediating quinine resistance

Of the 82 genes within the QN IC_90_ chromosome 7 QTL segment, 33 harbored nonsynonymous SNPs that were all present only in the QN-resistant Cam3.II parasite but not NF54. Within this segment, DMT1 was of particular interest as it is annotated as a putative drug/metabolite transporter (DMT) that, like PfCRT, is a member of the drug/metabolite exporter (DME) family within the DMT superfamily. This locus began ∼4 kb downstream of the chromosome 7 QN IC_90_ QTL peak (LOD = 10.4). DMT1 is predicted to be a 434-amino acid-long membrane transporter with 9 or 10 transmembrane domains based on TMHMM and TMpred analyses^44,45^, respectively (data not shown), whereas PfCRT has 424 amino acids and ten transmembrane domains. Cam3.II DMT1 harbors Y107N and S129L mutations that are not present in NF54. Our analysis of 2,512 *P. falciparum* genomes (in the Pf3k database), covering six Asian and eight African countries, revealed the frequent presence of Y107N in all countries, whereas S129L was only observed in Asia in samples that already harbored Y107N (**Extended Data 6a**).

Using ChimeraX MatchMaker, we calculated the structural similarity between the AlphaFold-predicted NF54 DMT1^46^ and cryo-EM derived 7G8 PfCRT (**Extended Data 7**)^47^. This comparison revealed similar folds in their respective transmembrane regions, with a good alignment of 1.27 Å between 47 pruned atom pairs that are predicted to be in the transmembrane domains. However, the overall alignment across all 300 atom pairs was poor (20.175 Å) most likely due to the flexible regions that are annotated by AlphaFold as “very low confidence”, where the Y107N and S129L mutations reside.

Sequence alignments with DMT1 orthologs in other *Plasmodium* species revealed strong conservation in the TMHMM-predicted transmembrane domains (**Supplemental Fig. 3a**). Phylogenetic trees (**Supplemental Fig. 3b**) revealed that *P. falciparum* DMT1 was the most similar to DMT1 orthologues in other laverania malaria species, consistent with *P. falciparum* originating from gorillas or other great apes^48^. In *P. falciparum*, *dmt1* was predicted to be potentially dispensable in cultured ABS parasites and to have a fitness cost (**Fig. 3a**) in a transposon mutagenesis screen^42^, although in the Pf3K genome database no isolates were found to have stop codons, suggesting essentiality *in vivo*.

### Mutant DMT1 may play a modulatory role in parasite susceptibility to quinine

To experimentally test whether DMT1 is a modulator of QN susceptibility *in vitro*, we applied CRISPR/Cas9-based gene editing (**Supplemental Fig. 2a**) to modify this locus at the Y107 and S129 positions in both parents. Editing was also applied to various progeny (H079, H030, H071, H062, and H046) that were selected to cover a range of QN susceptibilities and genotypes at the *dmt1*, *pfcrt* and *ftsh1* loci (**Table S4**). Editing was confirmed using PCR primers and Sanger sequencing (**Table S8**). We then conducted drug assays on these lines alongside their unedited parasites (**Fig. 5a** and **Table S10**). A statistically significant sensitization to QN was observed between the SNP-edited lines and their unedited parasites only for H079 (mutant for *pfcrt*, *dmt1* and *ftsh1*). There was, however, a trend of the Cam3.II and H071 DMT1 revertants (*dmt1*^WT^) having a lower QN IC_50_ than the unedited strain (∼20% and 19% mean reductions, respectively). In the H062 and H046 progeny (both Dd2 PfCRT, WT DMT1), mutating DMT1 did not cause a shift in QN susceptibility. These data suggest that the genetic background, and the specific constellation of how individual modulators including DMT1 are coinherited, play key roles in QN response.

**Fig. 5.**
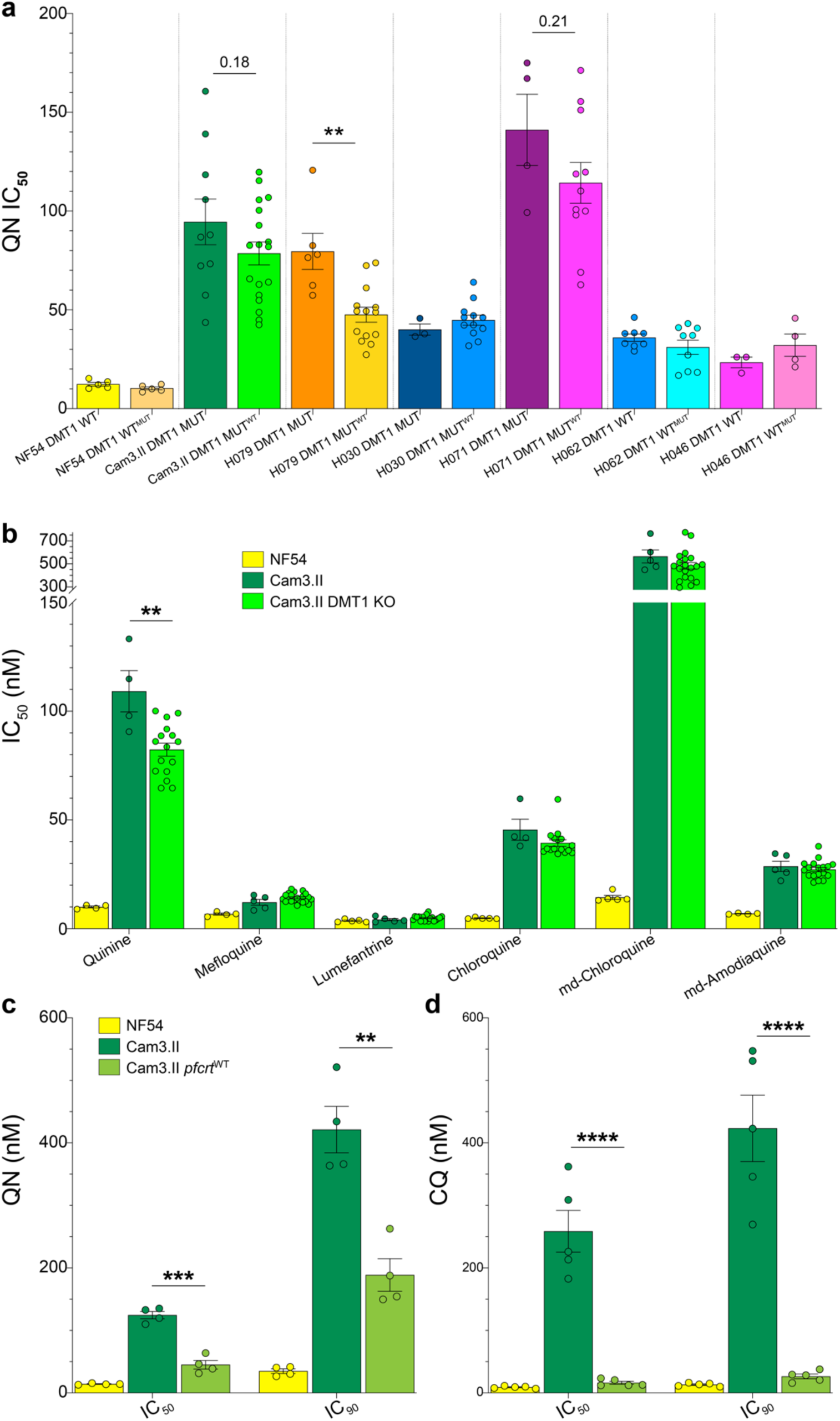
Cam3.II parasites lacking DMT1 or expressing WT PfCRT are significantly sensitized to QN. **a**, Quinine response of DMT1 SNP-edited parents and progeny as measured by mean ± SEM IC_50_ values. For each parasite, the parental unedited parasite (endogenous DMT1 haplotype) and the SNP-edited parasite (Y107N / S129L mutant or wild-type revertant haplotype) are indicated by dark and light colors, respectively. *p* values (Student’s *t*-test) are indicated (N,n=2−8,2). WT, wild-type; YNSL, Y107N / S129L mutation. **b**, Quinine, mefloquine, lumefantrine, chloroquine, md-Chloroquine, md-Amodiaquine response in Cam3.II DMT1 knockout parasites as measured by mean ± SEM IC_50_ values. *p*-values (Student’s *t*-test) are indicated for the Cam3.II KO strain vs the Cam3.II parent (N=4−21, with technical duplicates). **c-d**, Quinine (**c**) and chloroquine (**d**) response in Cam3.II *pfcrt*^WT^ revertant parasites as measured by mean ± SEM IC_50_ and IC_90_ values. *p*-values (Student’s *t*-test) are indicated for the Cam3.II *pfcrt*^WT^ strain vs the Cam3.II parent (N=4−5, with technical duplicates). \**p* < 0.05; \*\**p* < 0.01; \*\*\**p* < 0.001; \*\*\*\**p* < 0.0001.

We also profiled DMT1 SNP-edited lines and unedited isogenic controls against CQ, md-CQ, and the related 4-aminoquinoline md-amodiaquine (md-ADQ) (**Supplemental Fig. 4a-c** and **Table S10**). No statistically significant differences were observed, except for the H071 isogenic pair. H071 DMT1 revertants became less resistant to CQ and md-CQ than their unedited, CQ-resistant isogenic line, possibly due to an altered interaction with another gene in this recombinant background. The Cam3.II DMT1 revertant parasites did not become less resistant to md-CQ, CQ, and md-ADQ, compared with the isogenic DMT1 mutant lines. These results suggest that the modulatory role of DMT1 is mostly QN-specific.

### Phenotypic profiling of DMT1 knockout strains validates DMT1 as a marker of quinine resistance

To further investigate the role of DMT1, we generated KO parasites in Cam3.II parasites (**Supplemental Fig. 5**). Phenotypic characterization of four DMT1 KO clones, plus parental Cam3.II and NF54 parasites, showed that the DMT1 KO led to 25–30% lower QN IC_50_ and IC_90_ levels in the Cam3.II background (**Fig. 5b** and **Supplemental Fig. 4d** and **Table S10**). These data are consistent with DMT1 contributing to a multifactorial basis of QN resistance. KO parasites showed increased susceptibility to MFQ, LMF, and CQ at the IC_90_ but not IC_50_ level, evoking earlier findings of overlapping mechanisms of parasite susceptibility to these structurally partially related drugs (**Extended Data 1a)**^8,9,49,50^.

### Gene editing confirms a contribution for mutant PfCRT in QN resistance

Our QTL analysis with QN IC_50_ values revealed a chromosome 7 segment that also included *pfcrt*. To test this association, we reverted the *pfcrt* Dd2 allele in Cam3.II parasites to the WT (NF54) allele, using an established zinc finger nuclease-based gene editing method^51^. Dose-response assays showed a significant reduction in the QN IC_50_, and to a lesser extent QN IC_90_, values in the Cam3.II genetic background (**Fig. 5c**). Values remained higher than the reference NF54 line, consistent with *pfcrt* being only one component of a multigenic basis of QN resistance. Reversion of mutant *pfcrt* to the WT allele caused Cam3.II parasites to revert to a fully CQ-sensitive phenotype (evident at both IC_50_ and IC_90_ levels; **Fig. 5d**), as expected given the primary role for this gene in mediating CQ resistance.

### DMT1 localizes to the digestive vacuole, lipid bodies, parasitophorous vacuolar membrane, as well as structures associated with vesicular trafficking

To explore the subcellular localization of this novel transporter, we generated DMT1-3×HA-tagged parasites in NF54 parasites (**Supplemental Fig. 6**). Widefield deconvolution confocal microscopy with anti-3×HA antibodies showed a punctate pattern evident in all ABS parasites. We then co-stained parasites with antibodies or dyes specific for specific organelles or subcellular compartments (**Fig. 6a-g** and **Extended Data 8a-i**) and quantified their colocalization with DMT1-3×HA (**Table S11** and **Extended Data 8j**).

**Fig. 6.**
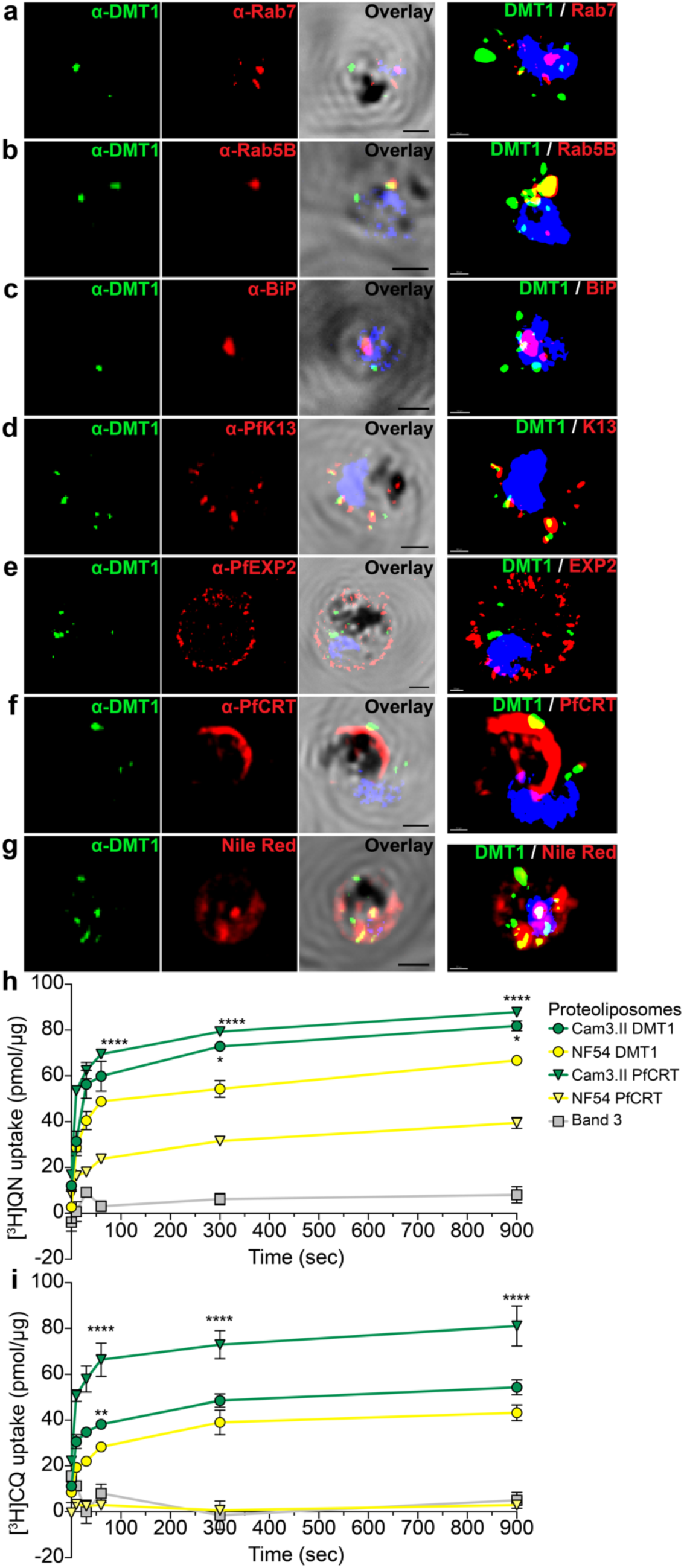
DMT1 shows localization to structures associated with vesicular trafficking as well as the digestive vacuole and exhibits differential uptake of [^3^H]QN between the NF54 and Cam3.II isoforms. **a**-**g**, Representative immunofluorescence assays (IFA) showing DMT1-3×HA-tagged parasites stained with antibodies or dyes: α-HA (DMT1, green), DAPI (nuclear, blue). **a**, α-Rab7 (late endosome), **b**, α-Rab5B (early endosome), **c**, α-BiP (ER), **d**, α-PfK13 (ER, vesicles, near cytostome), **e**, α-PfEXP2 (parasitophorous vacuolar membrane), **f**, α-PfCRT (digestive vacuole membrane), or **g**, Nile Red (lipid bodies). IFA images are shown as 3D volume reconstructions. Scale bars: 1 μm for IFA (black), 0.5 μm for 3D volume reconstructions (white). **h**,**i**, Time course of 300 nM [^3^H]QN and 300 nM [^3^H]CQ uptake measured with PfCRT (NF54 or Cam3.II), DMT1 (NF54 or Cam3.II), or Band 3-containing proteoliposomes. Data are mean ± SEM of *n* = 3 independent experiments with technical duplicates. Statistical significance between NF54 and Cam3.II variants for each protein was determined by unpaired Student’s *t*-tests corrected for multiple testing (**h**, **i**). \**p* < 0.05, \*\**p* < 0.01, \*\*\*\**p* < 0.0001.

We observed partial colocalization of DMT1 with the DV-resident markers PfCRT and plasmepsin 2 (PfPM2), with mean ± SEM overlap of 29.2 ± 8.0% and 16.7 ± 2.5%, respectively (**Fig. 6f** and **Extended Data 8f,g**). However, the mean colocalization values were not as high for some other organelles, indicating that DMT1 only partially overlapped with the DV membrane. In some parasites DMT1 almost fully (>80%) colocalized with lipid bodies (stained using Nile Red; 43.6 ± 10.8%) (**Fig. 6g** and **Extended Data 8h**), the endoplasmic reticulum (ER) (39.8 ± 10.3%) (**Fig. 6c** and **Extended Data 8d**), and the parasitophorous vacuolar membrane (PVM) (40.9 ± 12.4%) (**Fig. 6e** and **Extended Data 8e**). In other parasites, however, there was minimal overlap, leading to moderate colocalization values (shown in parentheses when averaged between these two patterns). This discrepancy may be due to DMT1 being trafficked within the parasite. Interestingly, Nile Red has been shown to localize to DV-adjacent small cytoplasmic neutral lipid droplets that contain di- and triacylglycerols in *P. falciparum*, which may explain why we observed colocalization with both Nile Red and PfCRT^52–54^. DMT1 also partially colocalized with Rab5B-positive early endosomes (27.9 ± 7.2%) (**Fig. 6b** and **Extended Data 8c**), and less so with Rab7-positive late endosomes (10.2 ± 5.7%) (**Fig. 6a**) and Rab11A-positive post-Golgi compartments (10.8 ± 4.5%) (**Extended Data 8i**). These observations suggest a role for DMT1 in vesicular trafficking. DMT1 also showed some colocalization with K13, the ART resistance mediator that is present in cytostomes at the plasma membrane, the ER, and internal vesicular structures^55–57^. DMT1 did not colocalize with the mitochondria or the apicoplast.

### NF54 and Cam3.II DMT1 isoforms differentially transport [^3^H]QN

To determine whether DMT1 can transport QN, we purified and reconstituted NF54 (WT) and Cam3.II (Y107N / S129L) DMT1 isoform proteins in proteoliposomes^47^ and measured [^3^H]QN and [^3^H]CQ uptake (**Fig. 6h,i**). DMT1 isoforms were found to transport both [^3^H]QN and [^3^H]CQ, with the mutant Cam3.II isoform showing significantly higher [^3^H]QN uptake. These data provide a possible mechanistic explanation that links DMT1 as a potential transporter of QN and suggest that the QN mechanism of action and resistance may, in part, be mediated by mutant DMT1-mediated QN transport. For [^3^H]CQ, the highest level of transport was observed with Cam3.II PfCRT, contrasting with zero detectable transport with NF54 PfCRT. Lower [^3^H]CQ transport levels were observed with both the Cam3.II and NF54 DMT1 isoforms, consistent with mutant DMT1 being a factor only for QN and not for CQ resistance. Our data agree with previous studies partially associating mutant PfCRT with QN resistance, and observations of PfCRT transporting non-CQ drugs, thereby impacting the parasite drug response by potentially changing the drug concentration at their site of action^58^.

### QN partially inhibits heme detoxification

To evaluate the involvement of QN in heme detoxification, we tested its ability to inhibit conversion of heme to β-hematin (synthetic hemozoin) in a cell-free heme detoxification assay^59,60^. When compared with the known β-hematin inhibitors CQ (mean ± SEM IC_50_ = 11.2 ± 0.5 μM) and md-ADQ (mean ± SEM IC_50_ = 5.2 ± 1.0 μM), we observed a lower degree of inhibition with QN (mean ± SEM IC_50_ = 22.1 ± 1.2 μM) and MFQ (mean ± SEM IC_50_ = 33.8 ± 4.6 μM) (**Extended Data 1b** and **Table S12**). These results are consistent with the structural relatedness between QN and MFQ (**Table S13**) and prior evidence of partial involvement of these drugs in the heme detoxification pathway^61,62^.

## Discussion

Here, we report the identification of three genetic markers associated with a polygenic basis of *P. falciparum* ABS parasite resistance to QN, an antimalarial drug first used in the 1600s for which resistance has long been enigmatic. This was achieved using a genetic cross between QN-resistant Cam3.II and QN-sensitive NF54 parasites in a humanized mouse model, from which we recovered progeny representing 120 unique recombinant haplotypes. QTL mapping associated QN resistance with a dominant peak on chromosome 7 that included a previously uncharacterized drug-metabolite transporter, DMT1. A smaller upstream peak aligned with *pfcrt*, the primary determinant of resistance to the former gold standard drug, CQ. For both QN and CQ, we also observed a peak on chromosome 12, including *ftsh1*. Gene editing studies confirmed a role for *dmt1*, *pfcrt* and *ftsh1* in QN resistance, supporting these as candidate molecular markers for malaria molecular surveillance.

DMT1 is a predicted drug-metabolite exporter that localizes to the DV, lipid bodies, the PVM, and structures associated with vesicular trafficking, as evidenced by partial Rab5B and ER staining. Our results suggest that DMT1 may be trafficked by vesicular structures from the parasitophorous vacuolar membrane to either the DV membrane or DV-adjacent lipid bodies. With the latter, we speculate that DMT1 may be involved in lipid storage of phospholipids that are known to promote β-hematin formation^54,63^. Structurally, DMT1 resembles PfCRT, which resides solely on the membrane of the DV wherein heme accumulates and is detoxified via biomineralization of β-hematin dimers into inert hemozoin^26^. We were able to functionally link QN and DMT1 by showing that proteoliposomes loaded with purified recombinant DMT1 protein could mediate [^3^H]QN uptake, with the Cam3.II variant showing higher levels than the WT protein. These data suggest that mutant DMT1 might contribute to QN resistance via intracellular flux away from this drug’s site of action. We note that QN uptake was also observed with mutant PfCRT, more so than with WT PfCRT, suggesting an additional role for PfCRT in QN resistance. Those findings are consistent with gene editing-based evidence that parasites expressing WT PfCRT were more sensitive to QN than the isogenic line expressing Cam3.II PfCRT. The direction of transport is uncertain, as QN can at high concentrations inhibit hemozoin formation that occurs in the DV, yet this drug is also suspected to have additional, unknown targets located in the cytosol^19^.

DMT1 harbors two mutations (Y107N / S129L) in Cam3.II but not NF54. This gene was also WT in prior genetic cross parents, which would explain its prior lack of detection^16,29,38,64^. In cultured parasites, *dmt1* is nonessential and we found that Cam3.II DMT1 KO parasites were significantly sensitized to QN (by ∼25% at the IC_50_ level), without affecting parasite susceptibility to other antimalarials (CQ, md-CQ, md-ADQ, MFQ, and LMF). QN resistance, and the contribution of DMT1, remains complex. SNP-edited Cam3.II DMT1^WT^ parasites exhibited a ∼17% decreased QN susceptibility compared to their unedited parent, and a 20–40% decrease in two progeny genetic backgrounds, yet none in another. The larger reduction in QN IC_50_ levels in the Cam3.II KO line compared to the SNP-edited line suggests that the level of DMT1 protein expression may also be important for mediating QN resistance. An analogous finding applies to PfCRT, where reducing the protein expression level led to a corresponding reduction in CQ IC_50_ in CQ-resistant parasites^65,66^. Of note, we observed no significant shift in QN response when mutating DMT1 from WT to Y107N / S129L in three genetic backgrounds (NF54, H046 and H062), suggesting that *dmt1* is only one determinant of a multigenic basis of QN resistance that in a background-dependent manner also includes PfCRT.

Our QTL analyses identified an unexpected peak on chromosome 12 that overlapped for QN, CQ and md-CQ. Gene editing of prioritized candidates implicated *ftsh1* as a potential contributor to these resistance mechanisms, as the Cam3.II *ftsh1* MUT^WT^ parasites (where the D659G mutation was reverted to WT) became significantly more susceptible to all three drugs compared to unedited Cam3.II (**Fig. 4**). This gene encodes a membrane metalloprotease that is homologous to the bacterial membrane AAA+ metalloprotease^67^. In *P. falciparum* and *Toxoplasma gondii* (both Apicomplexa), FtsH1 is required for apicoplast biogenesis^39^. In *P. falciparum* D10 parasites, FtsH1 G489C (located in the peptidase domain) confers resistance to actinonin, an antibiotic that kills *P. falciparum* within a single parasite generation – unlike apicoplast ribosome-targeting antibiotics that cause delayed death^41,68^. It remains to be tested whether QN might also interfere with apicoplast biogenesis.

We observed strong linkage disequilibrium between *ftsh1* and the chromosome 7 segment harboring both *pfcrt* and *dmt1*, suggesting functional complementarity that might offset any physiological defects resulting from mutations in *pfcrt* or *dmt1*. A prime candidate is the excess of short globin-derived peptides observed in mutant *pfcrt* parasites^66,69–71^. Metabolomic assays are planned to assess whether peptide levels return to a normal baseline in progeny harboring both chromosome 7 and 12 loci. When considering how FtsH1 might relate to DV transporters, we note that the apicoplast is essential for isoprenoid biosynthesis, which prenylates Rab5 proteins required for vesicle trafficking to the DV and normal DV development^72,73^.

Future studies are required to test for changing prevalences of mutant *dmt1*, *pfcrt*, and *ftsh1* in areas where QN is being used. This will become increasingly important if QN use expands in response to the increasing detection of *P. falciparum* strains across Africa that are partially resistant to artemisinin and the threat that they pose to effective treatment of severe malaria with artemisinin derivatives. Further studies are also required to edit these mutations across additional parasite strains, and further analysis of whether these mutant genes operate epistatically as a complex genetic trait of resistance.

Lastly, our QN studies highlight the value of treating malarial infections with compounds having complex modes of action that require resistance to be multigenic. Despite QN being the oldest antimalarial drug, it remains broadly effective to this day. In future drug discovery efforts, chemical compounds that have multiple *P. falciparum*-specific targets and/or multiple low-level resistance modulators merit prioritization for clinical use.

## Methods

### Parasite *in vitro* culture

*P. falciparum* ABS parasites were cultured in human RBCs at 3% hematocrit, using RPMI 1640 supplemented with 25 mM HEPES, 2.1 g/L sodium bicarbonate, 10 mg/mL gentamicin, 50 mM hypoxanthine, and 0.5% (w/v) AlbuMAX II (Thermo Fisher Scientific). Parasites were maintained at 37°C under 5% O_2_/ 5% CO_2_/ 90% N_2_ gas conditions. RBCs and human serum were procured from the Interstate Blood Bank (Memphis, TN) and RBCs were also procured from the New York Blood Center (New York, NY).

### Genomic DNA (gDNA) extraction

Parasite genomic DNA was obtained from cultures at 5 to 10% parasitemia with mostly trophozoite and schizont stages. RBCs were lysed with 0.1% saponin in 1×PBS and parasite DNA extracted with the QIAamp DNA Blood Midi Kit (Qiagen) using a combined RNase and Proteinase K treatment. DNA concentrations were determined using a NanoDrop and/or the Qubit dsDNA BR assay kit (Thermo Fisher Scientific).

### Parasite drug-susceptibility assays

Parasites were synchronized for at least two consecutive ABS cycles prior to starting drug susceptibility assays and at least once a week thereafter, by exposing ABS cultures to 5% D-sorbitol (Sigma) for 15–20 min at 37°C to remove mature parasites, followed by a wash step. In drug susceptibility assays, predominantly ring-stage parasites were exposed to a serially diluted range of drug concentrations (twofold, 10-point dilutions for all drugs except for QN that used 12-point dilutions), or 700 nM DHA (kill control), or a no drug control. Assays started at 0.2% parasitemia and 1% hematocrit in 96-well flat-bottom plates incubated at 37°C for 72 h in 5% O_2_ / 5% CO_2_ / 90% N_2_. Final parasitemias were measured using an IntelliCyt iQue Screener Plus cytometer (Sartorius), a BD Accuri C6 Plus cytometer with a HyperCyt plate sampling attachment (IntelliCyt), or a BD FACSCelesta flow cytometer with the BD High Throughput Sampler (HTS). Cells were stained with 1×SYBR Green I (Invitrogen) and 100 nM MitoTracker DeepRed FM (Invitrogen) for at least 20 min and diluted in 1×PBS prior to sampling (typically ∼8,000-10,000 RBCs per well). Parasitemia in each well was determined as the percentage of MitoTracker-positive SYBR Green I-positive infected RBCs within the cell subpopulation gated in the FSC vs SSC plot. Parasitemia was then normalized by subtracting the background parasitemia in the 700 nM DHA kill control, and 50% inhibitory concentration (IC_50_) and 90% inhibitory concentration (IC_90_) values were calculated by nonlinear regression analysis (Prism version 10, GraphPad). For that analysis, we used a 4-parameter curve fitting where minimum and maximum parasitemias were set to 0% and 100% growth, respectively. Drug susceptibility assays were performed with QN, CQ, md-CQ, md-ADQ, LMF, and MFQ. Statistical significance was determined by nonparametric, two-tailed Mann-Whitney *U*-tests for QN, CQ, and md-CQ QTLs and Student’s *t*-tests for all other drug assays (Prism version 10, GraphPad).

### Genetic cross

NF54 and Cam3.II *P. falciparum* gametocytes were generated as previously described^33^. Briefly, parasites were propagated in RPMI 1640 media containing 10% (v/v) human serum, at 4% hematocrit and 37°C in a glass candle jar. Cultures were initiated at 0.5% asynchronous asexual parasitemia from a low-passage stock and maintained up to day 18 with daily media changes but without any addition of fresh RBCs. Day 15 to 18 cultures, containing largely mature gametocytes, were used for mosquito feeds. Cultures were centrifuged (108 × g, 4 min) and the parasite pellet was adjusted to 0.3% gametocytemia in a mixture of human O+ RBCs supplemented with 50% (v/v) human serum. To conduct a genetic cross between NF54 and Cam3.II, we used a 1:1 ratio of gametocytes at a final gametocytemia of 0.3%. Gametocytes were fed for 1 h to 4–6-day old *Anopheles stephensi* mosquitoes (Liston strain) that had been starved overnight, using glass membrane feeders. Unfed mosquitoes were removed after feeding. Fed mosquitoes were then maintained on a 10% sugar solution at 25°C and 80% humidity with a 14:10 h (light:dark) cycle including a 2 h dawn/dusk transition. Human RBCs used to set up the cultures were collected weekly from healthy volunteers, with informed consent, under a Johns Hopkins University Institutional Review Board approved protocol (number NA_00019050). All experiments were performed in accordance with institutional guidelines and regulations.

On day 7 post-infected blood meal, mosquito midguts were harvested and stained with mercurochrome. Oocysts were counted by brightfield microscopy using a Nikon E600 microscope with a PlanApo 10× objective. We observed a ∼85% prevalence of infection, with an average of ∼20 oocysts per mosquito. Between days 14 to 16, salivary glands were harvested from ∼35 mosquitoes, homogenized and the sporozoites counted using a hemocytometer. On average we obtained 40,000 salivary gland sporozoites per mosquito.

All animal experiments were performed in accordance with the Animal Care and Use Committee (ACUC) guidelines and approved by the Johns Hopkins ACUC (Protocol M017H325), with modifications to a protocol reported previously^74^. FRG NOD human liver-chimeric (huHep) mice^75^ were purchased and shipped from Yecuris Corporation. Upon arrival we began withdrawal of the mice from 2-(2-nitro-4-trifluoromethylbenzoyl)-1,3-cyclohexanedione (NTBC). Mice were housed in a sterile facility with sterile bedding, food, and water. Mice were weighed daily and those that showed weight loss >10% (compared to pre-shipment weight) were treated with 300 μL of sterile saline via intraperitoneal injection and given a nutritional supplement in water (STAT; https://www.prnpharmacal.com/products/nutritional-supplements/stat/). Prior to infection, mice were allowed to recover from shipping for 2 to 3 days. Infection with *P. falciparum* sporozoites was performed either by mosquito bite or by intravenous (IV) inoculation. For the mosquito bite approach, mosquitoes were allowed to probe/feed for 17 to 19 min on mice anaesthetized with 150 μL ketamine/xylazine. Two mice (A and B) each received mosquito bites from approximately 60 mosquitoes. Two other mice (C and D) were given prophylactic penicillin/streptomycin antibiotics and then IV-inoculated with 200,000 sporozoites dissected from mosquito salivary glands. Mice infected by mosquito bite remained stable and healthy appearing for the remainder of the experiment. In contrast, both mice infected by IV inoculation lost ∼5% of their body weight relative to their pre-shipment weight and were given extra rehydration gel food. On days 5.5 and 6.5 post-sporozoite infection, 450–500 μL of RPMI-washed human RBCs at ∼70% hematocrit were IV-inoculated into each mouse (except for an intraperitoneal injection for Mouse B due to scar tissue at the tail vein injection site). Mice were given new dextrose, cages, and rehydration gel food on both days. At day 7 post-infection, ABS parasites were detected at 0.3–0.8% parasitemia and blood was recovered by cardiac puncture of mice anesthetized with ketamine/xylazine (**Table S1**). Bulk freezes were made after one and two cycles in culture. The bulk freezes made after one cycle were thawed from each mouse and used to obtain progeny clones by limiting dilution cloning (referred to as clones from no drug pressure prior to cloning). The post-two cycle freezes for mice B and C were later thawed and pressured at 50 nM or 75 nM CQ, or 75 nM or 95 nM QN, for 3 days in 5 mL cultures at 3% hematocrit. Surviving parasites were then cloned and cryopreserved. Mouse B ‘no drug’, as well as mouse B and C QN 95 nM pre-selected bulks, were thawed and pressured at QN 0 nM, 140 nM (and later 240 nM), or 180 nM (and later 240 nM) for 4 days for bulk selection experiments. Drug-pressured bulks were cloned by limiting dilution and cryopreserved.

### Parasite strains and progeny cloning

The Cam3.II G8 parasite, shortened throughout as Cam3.II, is a clone of the Cam3.II parasite (also known as RF967 or PH0306-C, kindly provided by Dr. Rick Fairhurst, then at the NIH). NF54 (JHU stock) and Cam3.II parents and their genetic cross progeny were cultured in human RBCs as described above, with the addition of 10% (v/v) human serum. Parasites were maintained at 37°C under 5% O_2_/ 5% CO_2_ / 90% N_2_ gas conditions. After cloning by limiting dilution, parasites were adapted to media devoid of 10% human serum, and all phenotyping was conducted in serum-free media.

### Whole-genome sequencing (WGS) of genetic cross progeny pools and clones

Parasite samples were subjected to WGS using the Illumina Nextera DNA Flex library preparation protocol. Briefly, 150 ng of gDNA was fragmented and tagmented using bead-linked transposons and subsequently amplified by five cycles of PCR to add dual-index adapter sequences to the DNA fragments. The libraries were quantified, pooled, and sequenced on either an Illumina MiSeq using the MiSeq Reagent Kit v3 (paired-end 300 bp reads) or a NextSeq 550 system using the High Output Kit (paired-end 150 bp reads).

Progeny were sequenced either directly (for bulk pools and clones after QN 140 nM, 180 nM, and 240 nM pressure) or after clonal recombinants had been identified by microsatellite and SNP genotyping (for clones following no drug pressure or CQ 50 nM, CQ 75 nM, QN 75 nM, or QN 95 nM pressure). The WGS data for the parents and progeny clones are visualized in **Fig. 1b**. Sequence data were aligned to the *P. falciparum* 3D7 genome (PlasmoDB version 48) using Burrow-Wheeler Alignment (BWA)^76^. Reads that did not map to the reference genome and PCR duplicates were removed using SAMtools and Picard. The reads were realigned around indels using Genome Analyses Tool Kit (GATK) RealignerTargetCreator and base quality scores were recalibrated using GATK BaseRecalibrator.

Using SAMtools mpileup, variants were called and filtered based on quality scores (minimum base quality score ≥20, mapping quality >30, read depth ≥5) and multiallelic = FALSE to obtain high-quality SNPs that were annotated with SnpEff^77^. SNPs for highly polymorphic surface antigens and multi-gene families were removed. Homozygous SNPs that differed between the NF54 and Cam3.II parents were retained, defined as having > 90% alternate or reference allele frequency for NF54 or Cam3.II, respectively. We then selected one recombinant progeny per haplotype with the best sequencing coverage (lowest % of SNPs classified as heterozygous (allele frequency of >10% and <90%) or missing (no reads or reads that did not pass the variant caller’s quality filter, or DP<5)) to create a list of 120 haplotype genomes, alongside the parents. We further culled these loci if >90% of the 120 progeny had either NF54 or Cam3.II calls and were not multi-allelic. We also removed 48 SNPs that were classified as heterozygous or discrepant for NF54 and/or Cam3.II across multiple sequencing runs. This process led us to retain 13,116 high quality SNPs that differed between NF54 and Cam3.II.

Microsatellite sizes in the NF54 and Cam3.II parental strains were identified as described previously^78^. Briefly, to determine the confidence of a microsatellite based on the number of reads, a custom Python script utilizing the pysam module^79^ was written to call reads that harbored insertions or deletions at specified genomic loci within the respective windows. Integrative Genomics Viewer was used to verify the data.

### Identification of unique recombinants

PCR and gel electrophoresis microsatellite genotyping (n=11 markers; **Table S14**) was used to narrow down which of the 175 progeny obtained after no drug pressure were recombinant clones, before their gDNA was whole-genome sequenced. This work yielded 162 clonal progeny (listed in **Table S15** and **Table S16**), of which 71 were chosen for WGS analysis. To obtain additional unique recombinants, we also subjected bulk progeny pols from mice B and C to various CQ and QN drug pressures (**Extended Data 2b** and **Table S15**). We applied CQ 50 nM, CQ 75 nM, QN 75 nM, or QN 95 nM for 3 days, while we applied QN 140 nM (and later 240 nM), and QN 180 nM for 4 days on the ‘no drug’ bulks or the QN 95 nM pre-pressured bulks. After a drug wash-off and a recovery period, we obtained 133 clonal progeny from CQ pressure and an additional 292 from QN pressure. 296 out of 303 progeny obtained after CQ 50 nM, 75 nM and QN 75 nM or 95 nM selections were genotyped with nine markers: two sets of three microsatellite markers via multiplexed capillary electrophoresis-based genotyping (**Table S14**), as well as SNPs in *pfcrt* (A220S), *dhfr* (N51) and *pfmdr1* (Y184F). These 9 markers were used to screen for clonal recombinant progeny before a subset of 92 progeny were whole-genome sequenced. As the primary goal of the higher QN concentrations was to recover QN-resistant progeny, we split cloning plates 1:2 for progeny obtained after QN 140 nM, 180 nM, or 140+240 nM, and subjected plates to 72 h of 140 nM QN. Surviving resistant parasites then proceeded to WGS analysis for 95 out of 160 progeny. PCR and either fragment analysis or gel electrophoresis-based MS genotyping always correctly predicted the self-fertilized or recombinant status of the progeny, which was also confirmed by WGS.

To identify which progeny were isogenic and to assign unique haplotypes, WGS results for the 13,116 high quality SNP positions were hierarchically clustered for the parents and clonal progeny by average linkage using Gene Cluster 3.0^80^. The resulting dendrograms were visualized using Java TreeView^81^. The SNP inputs were as follows: NF54 allele = - 3, Cam3.II allele = 3, heterozygous or missing = 0. Clustering of the unique recombinant haplotypes and the two parents were also conducted. If isogenic progeny shared the same haplotype, then the sequence for the progeny with the best sequencing data (lowest percentage of mixed and missing SNP calls) was used as the representative haplotype for subsequent clustering analysis.

### Quinine bulk segregant analysis

Recombinant bulk pools from Mouse B and C (with or without prior QN 95 nM drug pressure) for the NF54×Cam3.II cross were used for QN BSA experiments (full sample list detailed in **Table S5**). Bulk pools were pressured for four consecutive days in a 5 mL culture at 3% hematocrit in culture media (as mentioned above) supplemented with 10% (v/v) human serum, starting at 1% total parasitemia. gDNA, used for library preparations and WGS, was harvested before (day 0 control) and after the 4-day selections (changing media every day with drug wash off after four days) at 0 nM, 140 nM, 180 nM, and 240 nM QN, and later (day 4+) when there was sufficient genomic material to harvest (∼2% parasitemia). The 0 nM no drug control well was set up at the same time as each drug selection, to ensure drug-specific killing of the pressured well and to harvest no drug control gDNA at the same timepoint as its corresponding drug-pressured well. This ‘no drug timepoint’ control was used to account for allele frequency changes due to *in vitro* culture alone.

The gDNA from of the 33 bulk pool samples (**Table S5**) was whole-genome sequenced and used for the QN bulk segregant analysis performed with the R package, QTLseqr^82^. As some WGS data from gDNA bulk cultures exhibited low coverage compared to the progeny clones, we used all 106,316 SNPs that distinguished the two parents. SNPs were then filtered using the filterSNPs function^82^ using the following restrictions: 0.1≤ reference allele frequency ≤0.9 to remove over- and under-represented SNPs in both bulks, (minimum total depth = 10) / (maximum total depth = 200) and minimum sample depth = 5 to filter extremely low and high coverage SNPs in both bulks and in each bulk separately, respectively. SNPs were also removed if there was ≥100 read depth difference between bulks. QTLs were identified by comparing QN-pressured bulks with control bulks using the QTL-seq and G’ approaches^83,84^. A window size of 100 kb was used (unless indicated otherwise for G’ analysis) to calculate the tricube-smoothed Δ(SNP-*index*) and G statistic (or G’) for each SNP within that widow. For the G’ analysis, outlier regions were filtered based on Hampel’s rule and QTLs were identified as regions with False Discovery Rate (Benjamini-Hochberg-adjusted *p*-values or *q*-values) <0.01. For the Δ(SNP-*index*) analysis, QTLs were identified as regions that pass the 95% and 99% confidence intervals.

### Progeny clone-based quantitative trait locus (QTL) mapping

The R package, R/qtl version 2^85^ was used to map QTL peaks (**Fig. 2** and **Extended Data 5b,c**), using progeny IC_50_ and IC_90_ values as “phenotypes” (**Fig. 1 c-e** and **Extended Data 5a** and **Table S4**) and their WGS data as “genotypes.” To identify significant QTLs for each drug response phenotype, 1,000 permutations of phenotypic data (IC_50_, IC_90_) were performed to obtain a distribution of maximum Log of the Odds (LOD) scores. These LOD scores were then used to calculate the LOD threshold at 95% probability. Fine mapping of the QTL segments was performed using Bayesian interval mapping at a 95% confidence level. Significant QTL segments are listed in **Fig. 2e**, and the genes within these segments are shown in **Table S7**.

### CRISPR/Cas9 plasmid construction

The DMT1 (PF3D7_0715800) Y107N/S129L mutation or WT with binding-site mutations was introduced into the cross parents (NF54, Cam3.II) and five progeny (B-QN95-140-A5, B-CQ75-1-H9, B4-5, B5-4, Clow6, C-QN95-180-F8) using an all-in-one CRISPR/Cas9 plasmid (pDC2-co*Sp*Cas9-h*dhfr*^86^). The *P. falciparum* codon-optimized Cas9 endonuclease was derived from *Streptococcus pyogenes* and was expressed under a *calmodulin* promoter. The plasmid also carried a human dihydrofolate reductase (h*dhfr*) selectable marker (conferring resistance to the antifolate WR99210) under a *P. chabaudi dhfr*-*ts* (*Pcdt*) promoter and a *U6* promoter for expressing the guide RNA (gRNA). The DMT1 WT and Y107N/S129L donors were synthesized in a pUC-GW-*amp* vector (Genewiz), with both donors also containing the same three silent (synonymous) mutations at the gRNA recognition site (referred to herein as binding-site mutations or bsmut). These mutations prioritized the PAM and/or the “seed” region <12 nt upstream of the PAM to prevent Cas9 endonuclease-mediated re-cleavage of the donor or the edited recombinant locus. Donors were PCR amplified from the pUC-GW-*amp* vector and subcloned by In-Fusion cloning into pDC2-co*Sp*Cas9-h*dhfr* at the *Eco*RI and *Aat*II restriction sites. A gRNA was selected using ChopChop version 3^87^ and Benchling (https://www.benchling.com), as close as possible to the mutation site, avoiding poly-T stretches of >3 Ts as they may cause premature transcription termination^88^, and prioritizing gRNAs with low off-target scores and at least 3 to 4 mismatches to other sites in the genome that are preferably in the PAM and/or the “seed” region. Double-stranded gRNA blocks were synthesized as single-stranded DNA (Eurofins Genomics) and then phosphorylated, annealed, and cloned into the *Bbs*I restriction sites of pDC2-co*Sp*Cas9-h*dhfr* by T4 DNA ligase cloning.

CRISPR/Cas9 was also used to generate Cam3.II parasites with WT ThzK (PF3D7_1239600) or the A301V mutant, WT FtsH1 (PF3D7_1239700) or the D695G mutant, and WT SAMC (PF3D7_1241600) or the I176K/S193A variant. Donors were synthesized in the pUC-GW-*amp* vector as described above with synonymous binding site (bsmut) mutations introduced for each gene for each of two separate gRNAs. The donors for ThzK and FtsH1 were subcloned into pDC2-co*Sp*Cas9-h*dhfr* as described above while the donor for SAMC was kept in the pUC-GW-*amp* vector and co-transfected instead due to the low complexity of *samc*. Two gRNAs per gene were cloned into the *Bbs*I restriction sites to generate two separate plasmids for each gene, as described above.

To generate a PfNF54 DMT1 3’ 3×human influenza hemagglutinin-based (HA) tagged parasite, a DMT1 donor with a homology region (HR) 1 (342 bp), recodonized 3’ end of DMT1 to disrupt homology with no stop codon (165 bp), spacer (6 bp), 3×HA tag with a stop codon, and 3’ untranslated region (UTR) HR2 (509 bp) was synthesized in a pUC-GW-*amp* vector (Genewiz), PCR amplified and cloned by In-Fusion cloning into the *Eco*RI/*Aat*II restriction sites of the pDC2-co*Sp*Cas9-h*dhfr* vector. Three gRNAs within the recodonized region were selected and cloned into the vector.

To generate a Cam3.II DMT1 KO using CRISPR/Cas9 gene editing, 1202 bp out of the 1305 bp-long native PfDMT1 exon was replaced with a selectable marker cassette expressing h*dhfr* under the control of the *PcDT* promoter and a 3’ UTR from *hrp2*. Human *dhfr* expression was selected using 2.5 nM WR99210. The left homology (499 bp) and right homology (437 bp) regions were PCR amplified from Cam3.II gDNA and inserted by In-Fusion cloning into the *Eco*RI/*Aat*II and *Apa*I restriction sites of the pDC2-co*Sp*Cas9-h*dhfr* vector, respectively. Three gRNAs within the deleted *dmt1* sequence were selected and cloned into the vector.

All final plasmids were purified using the NucleoBond Xtra Maxi kit (Macherey-Nagel) and confirmed by restriction digests and Sanger sequencing and are listed in **Table S9**. Primers used for cloning and verification are described in **Table S8**.

### Parasite transfections

ABS parasites were electroporated with purified circular plasmid DNA as described^89^. Per transfection, a 2.5 mL culture of 3% hematocrit, >5% ring-stage parasites was harvested and washed with 15 mL of 1× Cytomix. 75 μL of parasitized RBCs was then added to 50 μg of plasmid DNA and 1× Cytomix to a total volume of 440 μL, transferred to a 0.2 cm cuvette (Bio-Rad), and electroporated at a voltage of 0.31 kV and capacitance of 950 μF using a Gene-Pulser (Bio-Rad)^90^. For SAMC SNP editing, we cotransfected 50 μg each of pUC-GW with donor and CRISPR/Cas9 plasmid with one gRNA; for DMT1 KO, DMT1-3×HA, FtsH1 SNP, and ThzK SNP gene editing, we cotransfected 50 μg each of CRISPR/Cas9 all-in-one plasmids that contained the same donor but two distinct gRNAs. Electroporated cells were transferred from the cuvette to a well in a 6-well plate containing 3% hematocrit blood + media mixture, and media was generally changed ∼3–4 h post-transfection (or sometimes ∼15 h later). Starting the day after the transfections, the cultures were maintained in 2.5 nM WR99210 (if parasites have triple *dhfr* mutations; N51I/C59R/S108N) or 1 nM WR99210 (if WT *dhfr*) for six consecutive days^91^. WR9210 was procured from Jacobus Pharmaceuticals. Successful gene editing was assessed via Sanger sequencing of PCR products amplified directly from bulk cultures or from bulk culture gDNA. DMT1-3×HA-tagged parasite clones were obtained by limiting dilution. Successful *dmt1*-3×HA tagging was confirmed via PCR, Sanger sequencing, immunofluorescence, and Western blot assays. Successful *dmt1* KO was confirmed by PCR, Sanger sequencing, and gel electrophoresis. Oligonucleotide primers used in this study are listed in **Table S8**.

### Western blot confirmation of DMT1-3**×**HA tagging

Western blots were performed with lysates from NF54 DMT1-3×HA-tagged parasites and NF54 PRELI-3×HA-tagged parasites (positive control). Parasite cultures were washed twice in cold 1×PBS, and parasites were isolated by treatment with 0.05% saponin in PBS. Released parasites were lysed in 4% SDS, 0.5% Triton X-100 and 0.5% PBS supplemented with 1×protease inhibitors (Halt Protease and Phosphatase Inhibitor Cocktail, Thermo Scientific) and Pierce Universal Nuclease for Cell Lysis (Thermo Scientific). Samples were centrifuged at 14,000 rpm for 10 min to pellet cellular debris. 4×Laemmli Sample Buffer (Bio-Rad) was added to lysates and samples were denatured at 50°C for 15 min (without boiling to prevent aggregation of the DMT1 transmembrane protein). Proteins were electrophoresed on precast 4–20% Tris-Glycine gels (Bio-Rad) and transferred onto nitrocellulose membranes. Western blots were probed with a 1:1,000 dilution of primary antibodies to HA (mouse mAb; Cell Signaling), followed by a 1:5,000 dilution of anti-mouse IgG H&L HRP-conjugated secondary antibodies (Abcam). Western blots were revealed using Pierce ECL Western Blotting Substrate (Thermo Scientific) and imaged on a ChemiDoc system (Bio-Rad).

### Indirect immunofluorescence assays (IFAs)

Harvested NF54 DMT1-3×HA-parasitized RBCs were washed twice in 1×PBS and fixed in methanol-free 4% (v/v) formaldehyde (Pierce), supplemented with 0.0075% (v/v) glutaraldehyde (Electron Microscopy Sciences) in 1×PBS for 30 min at room temperature while stationary, followed by one 1×PBS wash. For mitochondria staining, harvested parasites were washed once with pre-warmed RPMI, stained with 50 nM MitoTracker Red CMXRos (Invitrogen) at 37°C in the incubator while stationary prior to the 1×PBS wash and fixation, and were kept covered throughout. Cell membranes were then permeabilized in 0.1% Triton X-100 in 1×PBS for 30 min at room temperature while stationary, followed by three 1×PBS washes. Autofluorescence was quenched using 0.1 M glycine in 1×PBS for 15 min at room temperature while rotating. Blocking was performed with 3% (w/v) bovine serum albumin (BSA) and 0.1% Tween 20 (v/v) in 1×PBS overnight at 4°C while rotating. Cells were incubated with primary antibodies for 2–4 h at room temperature while rotating, with dilutions ranging from 1:50 to 1:200, followed by a 1 h incubation with a species-specific fluorophore-conjugated secondary antibody diluted in 3% BSA and 0.1% Tween 20 in 1×PBS. As primary antibodies, we used rabbit anti-binding immunoglobin protein (BiP) (1:200) (kindly provided by Min Zhang), mouse anti-PfK13 (1:100) (clone E3^56^), rabbit anti-Rab11A, rat anti-Rab5B or -Rab7 (1:50) (kindly provided by Dr. Gordon Langsley), rabbit anti-PfACP (1:200) (kindly provided by Dr. Geoff McFadden), Nile Red (2.5 μg/mL) (Invitrogen), rabbit anti-PfEXP2 (1:200) (MR4), mouse anti-PfCRT (1:200) (clone 2^92^), and rabbit anti-plasmepsin 2 (PM2) (1:100) (BEI Resources). For primary antibodies raised in rabbits, we used Alexa Fluor Plus 594-conjugated goat anti-rabbit IgG (H+L) secondary (1:3000) (Invitrogen). For mouse primary antibodies, Alexa Fluor 594-conjugated goat anti-mouse secondary (1:2000) (Invitrogen) were used. For assays with rat primary antibodies, we used Alexa Fluor 594-conjugated goat anti-rat secondary (1:2000) (Invitrogen). Rat anti-HA (1:2000) (Millipore Sigma) and Alexa Fluor 488-conjugated goat anti-rat secondary (1:2000) (Invitrogen) antibodies were used to test for DMT1-3×HA expression. Samples were also costained with MitoTracker Red CMXRos, Nile Red, or primary antibodies raised in mice or rabbits. Mouse anti-HA (1:100) (Cell Signaling) and Alexa Fluor 488-conjugated goat anti-mouse secondary (1:1000) (Invitrogen) antibodies were used to probe for DMT1-3×HA in co-stains with primary antibodies raised in rats.

Thin blood smears of stained RBCs were prepared on microscope slides and mounted with high-performance/high-tolerance cover slips (Zeiss) using ProLong Diamond Antifade Mountant with DAPI (Invitrogen) pre-warmed to room temperature. Mounted samples were cured overnight and were imaged using a Nikon TiE Eclipse inverted microscope with a Nikon A1 scanning confocal with a GaAsP spectral detector, and a CFI Plan Apochromat Lambda oil immersion objective with 100× magnification (1.45 numerical aperture). 0.2 μm step size Z-stacks were taken for each parasitized RBC. NIS-Elements version 5.02 (Nikon) was used to control the microscope and camera and crop images. Imaris version 9 (Oxford Instruments) was used to perform deconvolution using 5 (for DAPI) or 10 (for HA tags and parasite proteins) iterations of the Richardson-Lucy algorithm for each image and quantify colocalization of the deconvolved Z-stacks using the Colocalization Module. Trophozoite-stage parasites were used to quantify colocalization of DMT1 with organelles, as there was more spatial distribution of organelles than in rings and trophozoites were less prone to variability than the multinucleated schizonts. Statistical analyses were performed with Prism version 10 (GraphPad). Fiji^93^ version 2.11.0 (ImageJ) was used to adjust brightness and contrast, overlay channels, and prepare montages. For 3D rendering of the deconvolved images, we used the Imaris software.

### Sequence alignment

The DMT1 protein sequence for the *P. falciparum* NF54 (PF3D7_0715800) strain and the orthologous DMT1 sequences from reference strains of *P. reichenowi* (PRCDC_0713100), *P. adleri* (PADL01_0713800), *P. gaboni* (PGSY75_0715800), *P. vivax*(PVP01_1424900), *P. knowlesi* (PKNH_1424800), *P. ovale* (PocGH01_14032100), *P. malariae*(PmUG01_14040800), *P. cynomolgi* (PcyM_1426000), *P. chabaudi* (PCHAS_1423900), *P. berghei* (PBANKA_1422100), *P. yoelii* (PY17X_1424100), and *P. gallinaceum* (PGAL8A_00520500) were obtained from the PlasmoDB.org database^94^. Clustal Omega^95^, which uses the HHalign algorithm, was used to perform multiple sequence alignments and generate phylogeny trees (**Supplemental Fig. 3**). EMBOSS Needle, which uses the Needleman-Wunsch algorithm, was used to perform global pairwise sequence alignments and calculate the percentage of positions in the alignment that are identical^96^. Both bioinformatic tools were accessed through EMBL-EBI^97^.

### DMT1 NF54 and Cam3.II protein expression and purification, and QN and CQ transport measurements

The NF54 (QN- and CQ-sensitive) and Cam3.II (QN- and CQ-resistant) variants of the DMT1 and PfCRT proteins as well as control Band 3 protein (anion transport control) were expressed, purified, and reconstituted in preformed liposomes as described previously^47^, with some modifications. The full-length NF54 and Cam3.II *pfdmt1* were cloned into pEG BacMam vector^98^ with the C-terminal tobacco etch virus protease cleavage site (ENLYFQSYV) and a decahistidine affinity tag followed by a streptavidin affinity tag (WSHPQFEK).

For protein expression, the NF54 and Cam3.II bacmid and baculovirus were generated using a BacMam method^98^. The baculoviruses were produced in Sf9 cells (Expression System) and added to HEK293 GnTi^-^ cells (ATCC, CRL-3022) incubated at 37°C for 20– 24 h in the presence of 5% CO_2_ and 60% humidity. We then added 10 mM sodium butyrate (Sigma-Aldrich) and further incubated at 37°C for 48 h before harvesting them. Cell pellets were homogenized in low-salt buffer (10 mM HEPES pH 7.5, 10 mM KCl, 10 mM MgCl_2_, 0.5 mM PMSF, cOmplete EDTA-free protease inhibitor cocktail, 10 μg/mL DNase I, and 8 μg/mL RNase) in a glass homogenizer. Membrane fractions were isolated by ultracentrifugation at 134,000 × g in a type 45 Ti rotor (Beckman Coulter). Membrane fractions were further homogenized and washed twice with high-salt buffer (10 mM HEPES pH 7.5, 10 mM KCl, 10 mM MgCl_2_, 1 M NaCl, 0.5 mM PMSF, cOmplete EDTA-free protease inhibitor cocktail, 10 μg/mL DNase I, and 8 μg/mL RNase) in a glass homogenizer followed by ultracentrifugation. The membrane fractions were homogenized and solubilized for 2 h in a buffer containing 20 mM HEPES pH 7.5, 200 mM NaCl, 0.5 mM PMSF, cOmplete EDTA-free protease inhibitor cocktail, and 1% n-dodecyl-β-d-maltopyranoside DDM/ 0.1% cholesteryl hemisuccinate (CHS). Insoluble material was removed by ultracentrifugation at 134,000 × g for 30 min. The supernatant was added to pre-equilibrated Ni^2+^-NTA resin (Qiagen) in the presence of 20 mM imidazole, and incubated at 4°C overnight with gentle rotation. The protein-bound resin was washed with 10 column volumes of buffer containing 20 mM HEPES pH 7.5, 200 mM NaCl, 60 mM imidazole, 0.1% DDM/ 0.01% CHS. The protein was eluted in buffer consisting of 20 mM HEPES pH 7.5, 200 mM NaCl, 200 mM imidazole, and 0.05% DDM/ 0.005% CHS. The eluted protein was further purified by loading on a Superdex 200 Increase 10/300 GL size-exclusion column (Cytiva Life Sciences) equilibrated with a buffer containing 20 mM HEPES pH 7.0, 150 mM NaCl, and 0.025% DDM/ 0.0025% CHS. Purified proteins were reconstituted in preformed liposomes made of total *E. coli* lipids:cholesteryl hemisuccinate 97:3 (w/w) at a protein-to-lipid ratio of 1:150 (w/w). The lumen of the proteoliposomes was composed of 100 mM KPi, pH 7.5, and 2 mM β-mercaptoethanol. Uptake of [^3^H]-CQ (300 nM, 1 Ci/mmol) or [^3^H]-QN (300 nM, 1 Ci/mmol) was measured after diluting PfCRT, DMT1, or Band 3-containing proteoliposomes in 50 μl of 100 mM Tris/MES, pH 5.5 in the presence or absence of the indicated compounds. Band 3 was used as a negative control, for which we expected no or minimal CQ and QN uptake. The protein-specific uptake (pmol/mg) was determined by subtracting the time-dependent accumulation of the tested compounds in control liposomes (lacking PfCRT, DMT1 or Band 3) from the accumulation measured in proteoliposomes containing one of these proteins. Data were collected in three independent experiments, performed in duplicate, and are represented as the mean ± SEM uptake at each timepoint.

#### β-hematin inhibition assays

A solution containing deionized H_2_O/ 305.5 mM lipophilic detergent NP-40 Surfact-Amps Detergent solution (ThermoFisher Scientific)/ dimethyl-sulfoxide (DMSO) was prepared at a v/v ratio of 70%/ 20%/ 10%, respectively, and 100 μL was added to columns 1–11 of a flat-bottomed 96-well plate. Working stocks of test and control compounds were constituted to 20 mM (5mM for md-ADQ), from which 20 μL (80 μL for md-ADQ) of each was added to wells in duplicate in column 12 together with deionized H_2_O (140 μL; 80 μL for md-ADQ) and 305.5 mM NP-40 detergent (40 μL). This effectively lowered the final drug concentration to 2 mM. Each compound (100 μL) was then serially diluted from columns 12 to 2 (column 1 served as a blank). A 25 mM hematin stock solution was prepared by sonicating hemin (Sigma Life Sciences) in DMSO for 3min and 178.8 μL of this solution was suspended in 20 mL of acetate buffer (1 M, pH 4.8) (made with sodium acetate (Sigma Aldrich) and acetic acid (Fisher Scientific)) and thoroughly mixed. The homogenous suspension (100 μL) was then added to all wells to give a final 0.5 M buffer and 100 mM hematin concentration, as well as a 0.5 mM drug concentration in column 12. Plates were covered and incubated at 37°C for 5 h after which 32 μL of a 50% pyridine solution (20% (v/v) H_2_O, 20% (v/v) acetone, 10% (v/v) 2 M HEPES buffer (pH 7.4) and 50% (v/v) pyridine (Sigma Aldrich)) was added to each well. 60 μL of acetone was then added to each well to assist with hematin dispersion. Unreacted hematin was quantified using the pyridine ferrihemochrome method^99^, which relies on aqueous pyridine specifically forming a low-spin complex with hematin but not β-hematin and the complex having an absorbance at 405 nm. The UV-vis absorbance of each well was read at 405 nm on a SpectraMax P340 plate reader. The β-hematin IC_50_ values for each compound were calculated from the absorbance values at 405 nm using sigmoidal dose-response curve fitting analysis (Prism version 10, GraphPad).

## Data and materials availability

All data needed to evaluate the conclusions in the paper are present in the paper and/or the Supplementary Materials. Raw reads of WGS data used in this study have been deposited in NCBI Sequenced Read Archive (SRA) under the BioProject accession number PRJNA1207213.

## Acknowledgments

Funding for this work was kindly provided by the National Institute of Allergy and Infectious Diseases, National Institutes of Health (R37 AI050234 and R01 AI185559 to D.A.F.; R01 AI147628 to F.M., M.Q. and D.A.F.; R01 AI132359 to P.S.) and the Bill & Melinda Gates Foundation (OPP1201387 to D.A.F. and P.S.). M.K. gratefully acknowledges funding support from the Japan Student Services Organization and a scholarship from the New York Hideyo Noguchi Memorial Society. S.M. gratefully acknowledges funding support Human Frontiers Science Program Long-Term Fellowship (LT000976/2016-L) and the NIH (R01 AI182318). M.S. was a recipient of a Johns Hopkins Provost’s Postdoctoral Fellowship. J.L.S-S gratefully acknowledges support from the NIH (K08AI163497) and from a Doris Duke Physician Scientist Award and a Louis V. Gerstner, Jr. Scholar Award. We also thank Bloomberg Philanthropies for their support of the Johns Hopkins Malaria Research Institute and the Insectary Core Facilities. The IFA studies used the resources of the Herbert Irving Comprehensive Cancer Center Confocal and Specialized Microscopy Shared Resource, funded in part through NIH/NCI Cancer Center Support Grant P30 CA013696, and with the assistance of Drs. Haojie Ji and Theresa Swayne. Brian Kloss and Renato Bruni of the New York Structural Biology Center kindly generated the DMT1 bacmids and viruses for the transport studies. We thank Susan Wyllie and Victor Corpas Lopez (University of Dundee, UK) for their kind help with this project. We dedicate this manuscript to Dr. Leila S. Ross, a brilliant scientist who recently passed away after a heroic battle with cancer.

---

**Extended Data 1.**
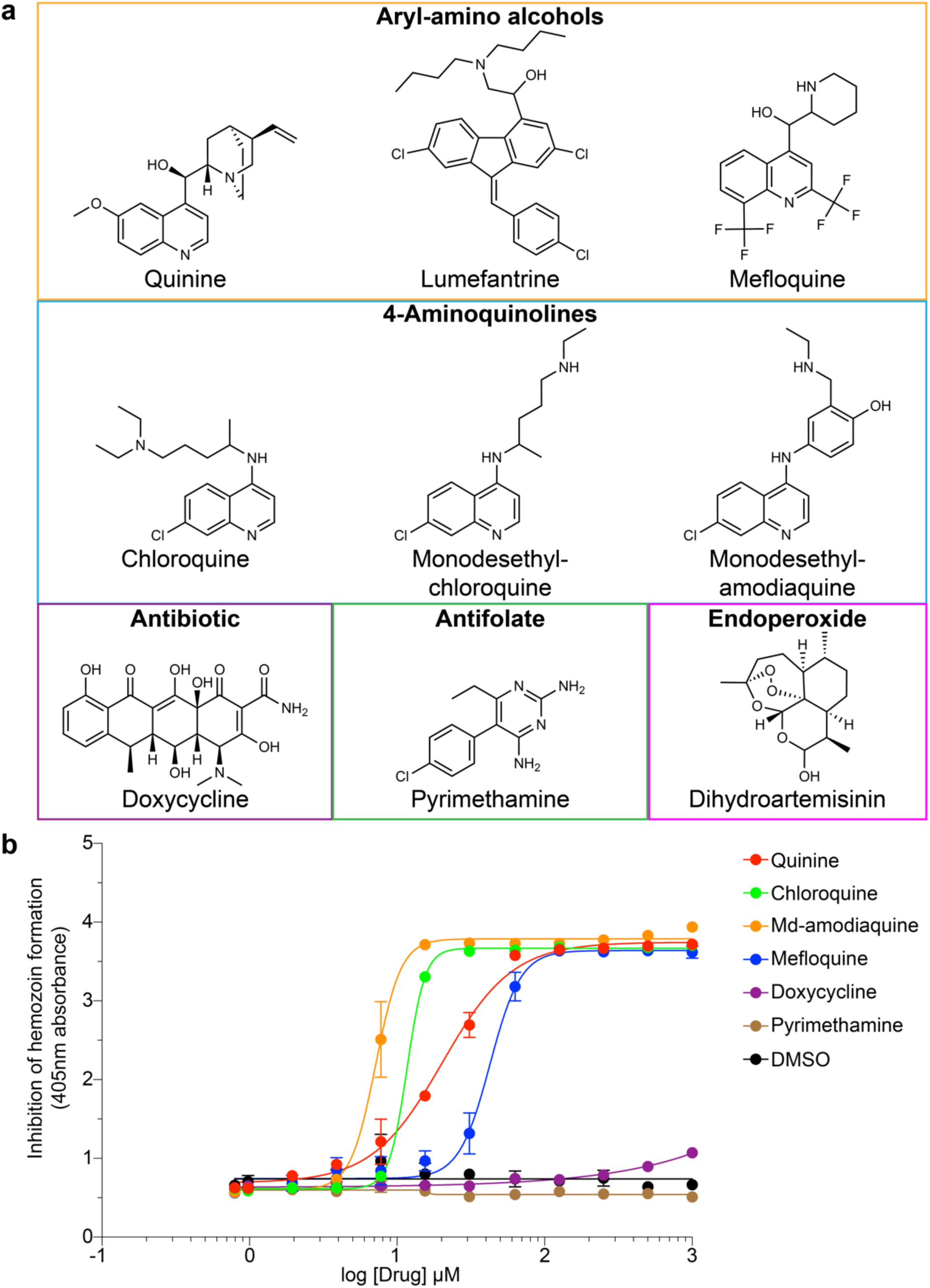
Quinine inhibits β-hematin formation to a lesser degree than chloroquine but more than mefloquine in a cell-free system. **a**, Chemical structures of the antiplasmodial compounds used in this study. Quinine, lumefantrine, and mefloquine are aryl-amino alcohols; chloroquine, monodesethyl-chloroquine, and monodesethyl-amodiaquine are 4-aminoquinolines; doxycycline is a tetracycline-class antibiotic; pyrimethamine is an antifolate; dihydroartemisinin is an active metabolite of artemisinin, a sesquiterpene lactone endoperoxide. Quinine and mefloquine have the aminoquinoline ring found in chloroquine, monodesethyl-chloroquine, and monodesethyl-amodiaquine. Chemical formulas, SMILES, and Tanimoto scores of these compounds are indicated in **Table S13**. **b**, Representative concentration-dependent inhibition of β-hematin formation by quinine, chloroquine, md-amodiaquine, mefloquine, two non-heme detoxification pathway-acting negative control antimalarials (doxycycline and pyrimethamine), and DMSO. Mean ± SEM IC_50_ values from N=3 experiments are listed in **Table S12**.

**Extended Data 2.**
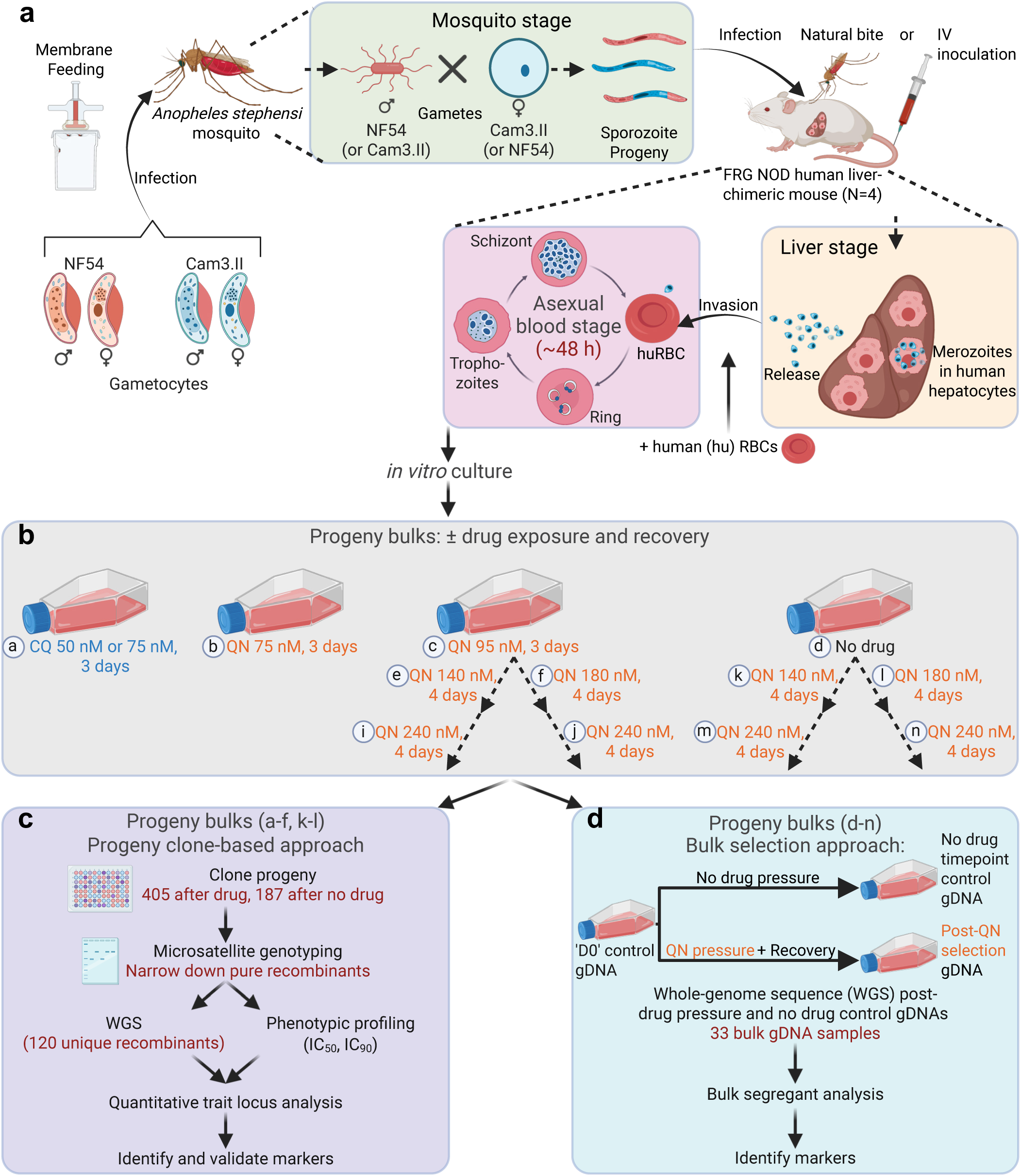
Workflow of the NF54×Cam3.II genetic cross, isolation of progeny, and bulk selection vs. progeny clone-based trait mapping. **a**, Overview of the genetic cross performed using *Anopheles stephensi* mosquitoes and FRG NOD human-liver chimeric mice (N=4). RBCs, red blood cells. **b**, Drug selection conditions were applied to progeny bulks, followed by a drug-free recovery period after which bulk genomic DNA was harvested and/or the progeny were cloned out. **c**,**d**, Genetic determinants of drug resistance were identified using progeny clone-based linkage approaches (**c**) or bulk selection approaches with bulk segregant analysis (**d**). Images were created with BioRender.com. The NF54 and Cam3.II parents are described in Fig. 1a, the mosquito and mouse infections are described in **Table S1**, and the total number of clonal progeny obtained are indicated in **Table S15**.

**Extended Data 3.**
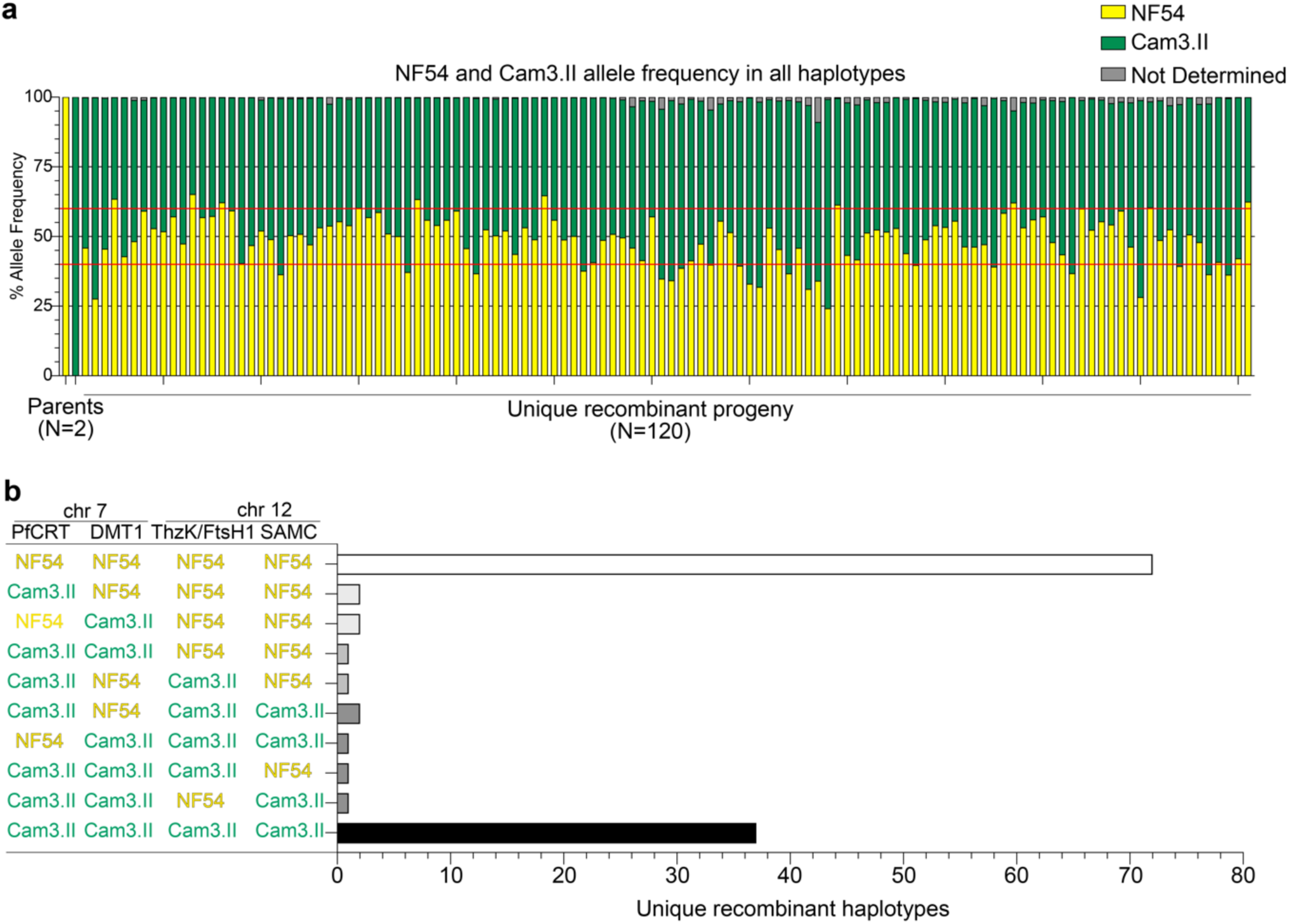
Genome-wide frequencies and *pfcrt*, *dmt1*, *thzk*/*ftsh1*, and *samc* allele representation in the recombinant progeny. **a**, Percentage of NF54 and Cam3.II allele frequencies averaged across the genome in 13,116 SNP positions for each of the parents and 120 recombinant progeny, ordered by haplotype number. **b**, Number of unique recombinant progeny with the wild-type NF54 or mutant Cam3.II alleles for *pfcrt, dmt1, thzk/ftsh1, and samc.* The mutant Cam3.II alleles for each gene are listed in Fig. 3a.

**Extended Data 4.**
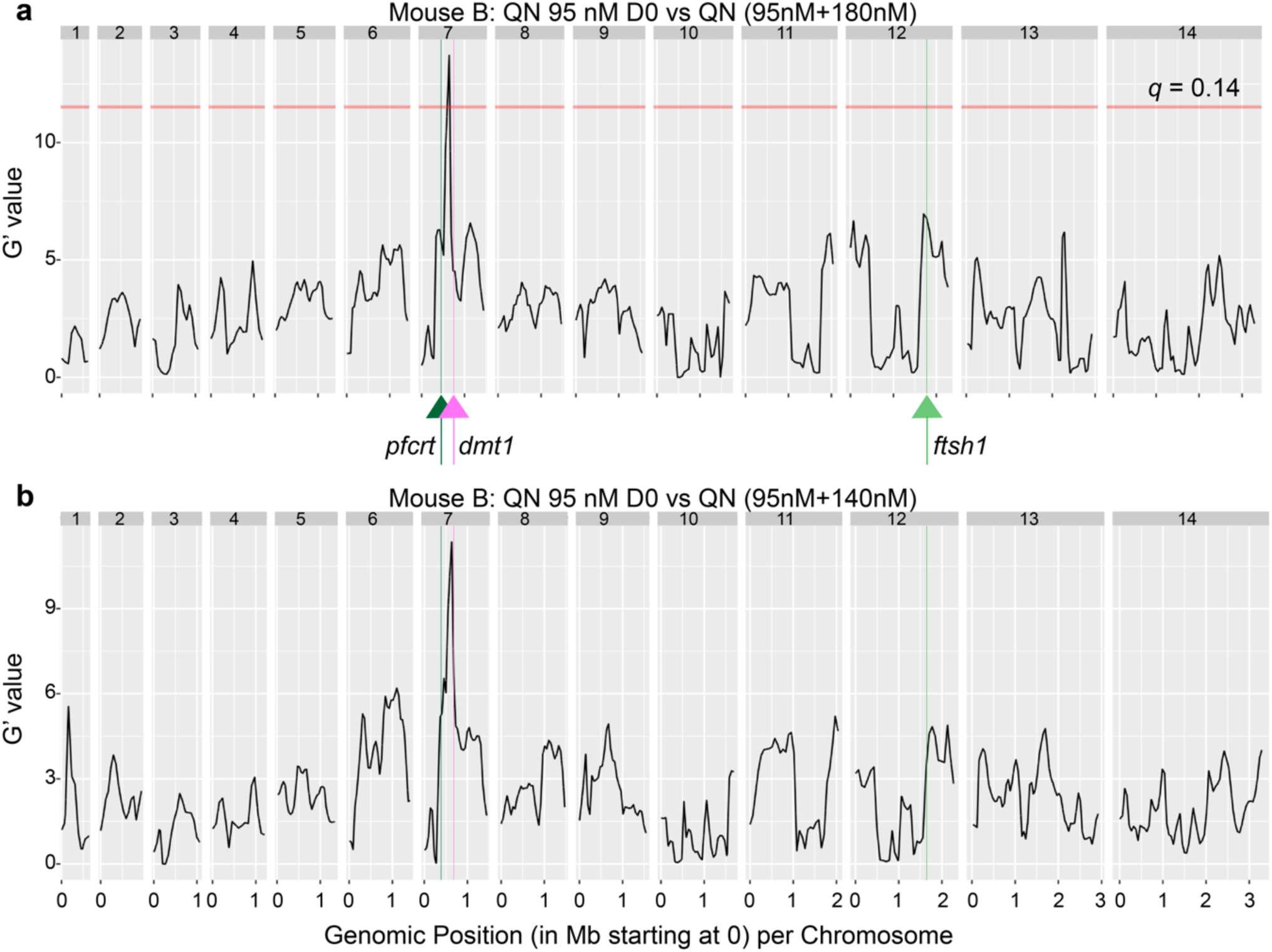
Bulk segregant analyses of progeny pools pressured with quinine or no-drug controls. **a**,**b**, Genetic loci enriched by quinine (QN) in drug-no drug pairwise comparisons. Black lines are G’ values comparing allele frequency between QN-pressured bulk progeny and control non-drug-pressured bulk progeny. The red line indicates the False Discovery Rate at the indicated threshold. Samples are described in **Table S5**. Quantitative trait locus regions are described in **Table S6**. Of note, mutant *pfcrt*/mutant *ftsh1, or* WT *pfcrt/*WT *ftsh1,* continued to be co-inherited in the QN-pressured progeny, as was seen in the non-drug-pressured progeny.

**Extended Data 5.**
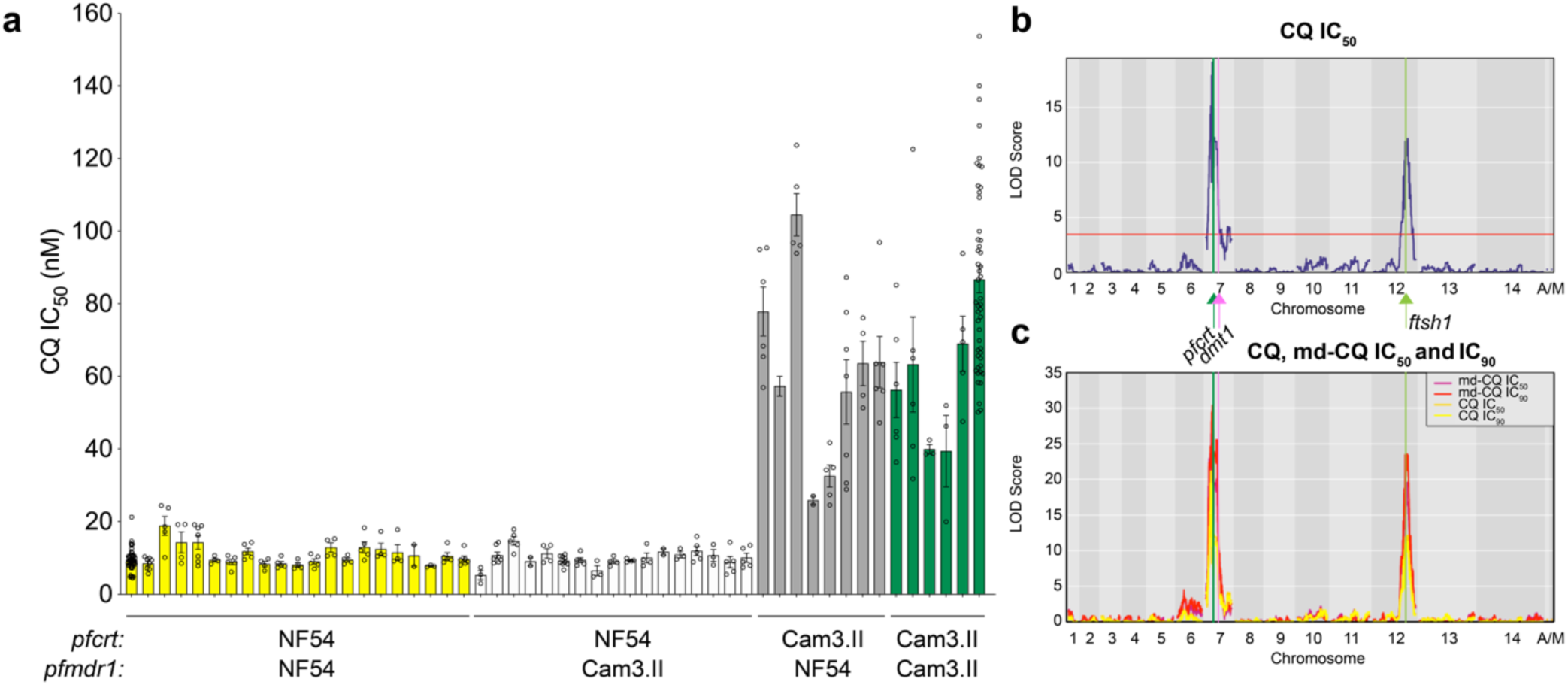
Phenotypic response of parents and recombinant progeny to chloroquine and monodesethyl-chloroquine closely align. **a**, Chloroquine (CQ) response as measured by mean ± SEM IC_50_ values. **b**, Logarithm of odds (LOD) plot for geometric mean CQ IC_50_ values. The red line indicates the 95% probability threshold. **c**, Overlay of monodesethyl-chloroquine (md-CQ) and CQ LOD plots. A/M, apicoplast/mitochondria (these are always coinherited). The highest LOD peaks for CQ and md-CQ align with *pfcrt* on chromosome 7 and the segment including *ftsh1* on chromosome 12. Note that *dmt1* was not within the dominant chromosome 7 peak for CQ or md-CQ, unlike for quinine.

**Extended Data 6.**
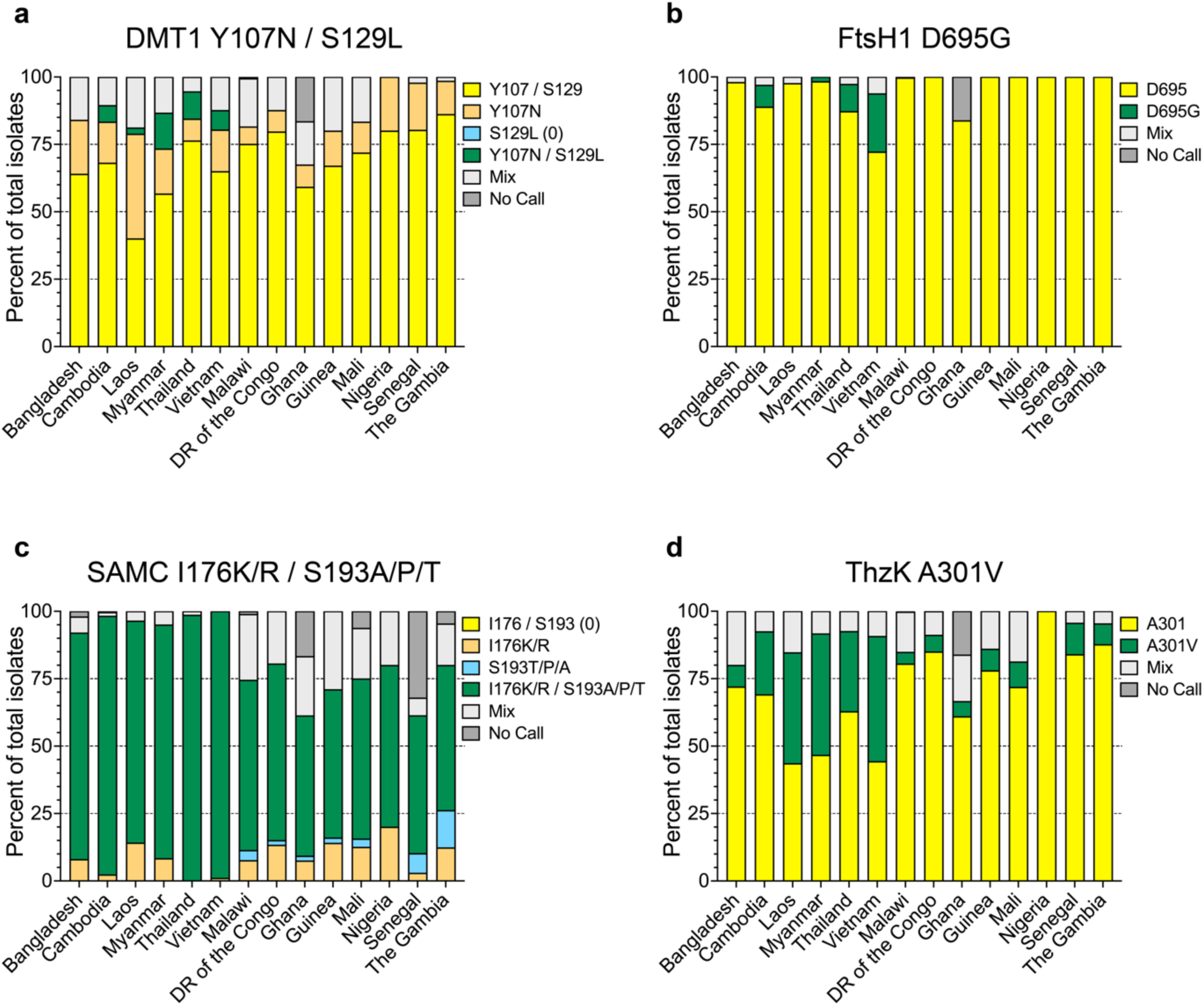
Clinical isolates harbor mutations in genes of interest within the chromosome 7 QN and chromosome 12 quinine (QN), chloroquine (CQ), and monodesethyl-CQ (md-CQ) quantitative trait locus segments. **a**-**d**, Frequency of the Cam3.II mutations in clinical isolates (Pf3k) of genes in chromosome 7 ((**a**), *dmt1:* PF3D7_0715800) and chromosome 12 ((**b**) *ftsh1:* PF3D7_1239700, (**c**) *samc:* PF3D7_1241600, (**d**) *thzk:* PF3D7_1239600)) loci, grouped by sampling country. Yellow indicates isolates with the NF54 wild-type haplotype covered by the indicated SNP positions; green, the Cam3.II mutant haplotype; light grey, mixed call; dark grey, genotype not called; orange or blue, single-amino acid mutants not corresponding to the double-mutant Cam3.II haplotype. (N=50 (Bangladesh), 570 (Cambodia), 85 (Laos), 60 (Myanmar), 148 (Thailand), 97 (Vietnam), 369 (Malawi), 113 (Democratic Republic of the Congo), 617 (Ghana), 100 (Guinea), 96 (Mali), 5 (Nigeria), 137 (Senegal), 65 (The Gambia); total=2,512 samples). These four top candidate genes are further described in Fig. 3a.

**Extended Data 7.**
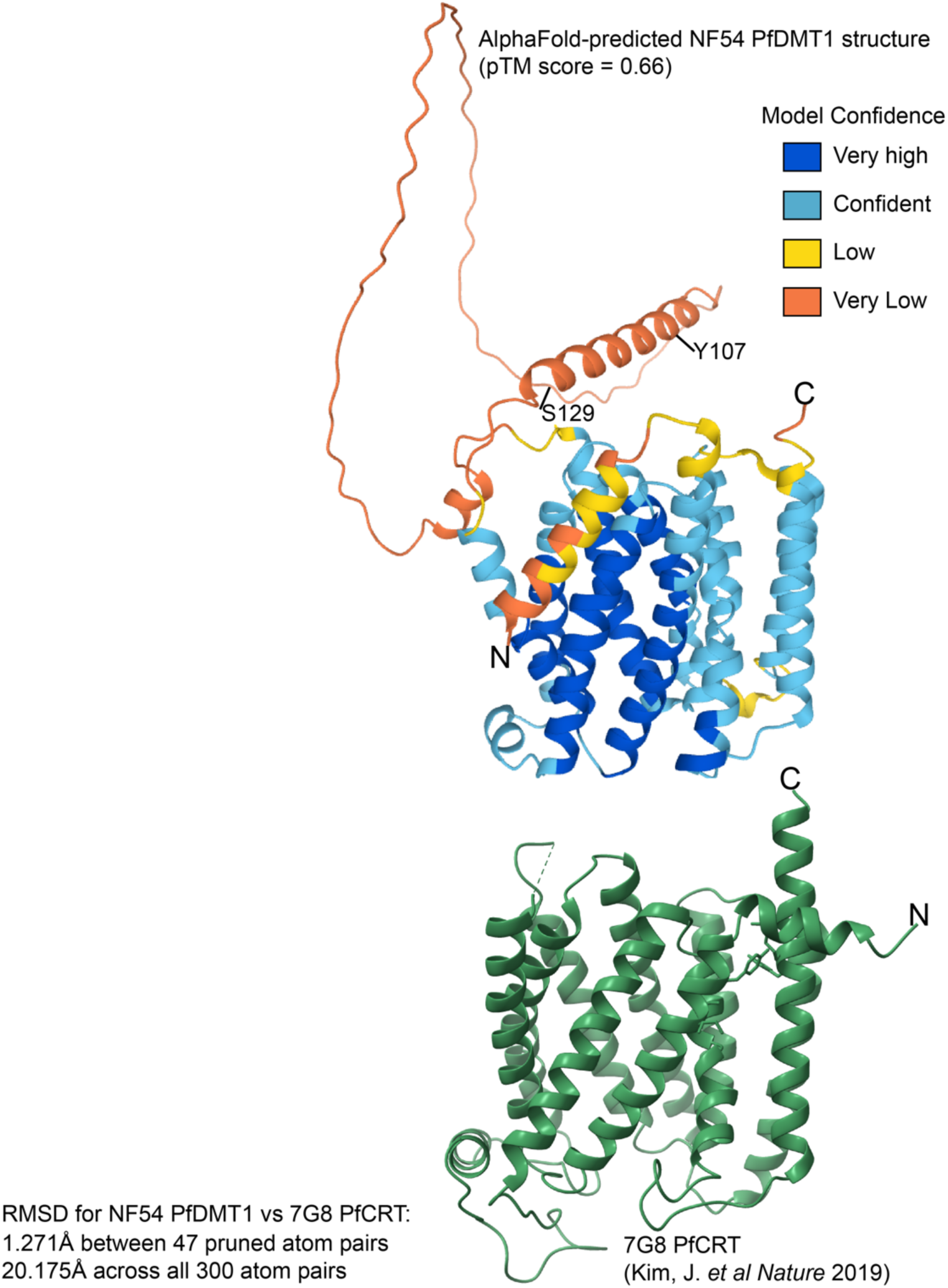
PfDMT1 is structurally similar to another drug metabolite transporter superfamily protein, PfCRT. Side-by-side comparison of the AlphaFold-predicted NF54 DMT1 structure and the cryo-EM-derived 7G8 PfCRT structure^47^. The predicted template modeling (pTM) is indicated for the AlphaFold NF54 DMT1 structure, which exceeds the 0.5 threshold for predicted fold reliability. The Y107 and S129 positions are indicated on the AlphaFold DMT1 structure. Root mean square deviation (RMSD) scores calculated in ChimeraX MatchMaker are indicated for PfCRT and DMT1.

**Extended Data 8.**
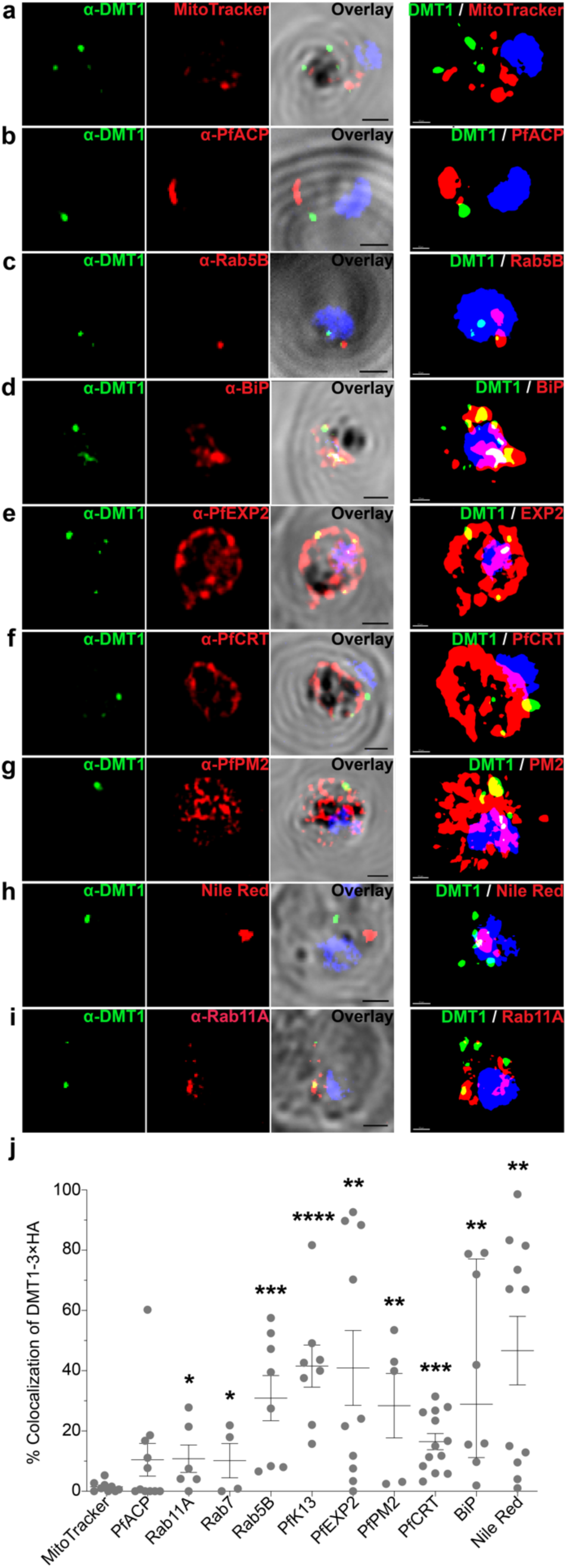
DMT1 shows localization to structures associated with vesicular trafficking as well as the digestive vacuole. **a**-**i**, Representative immunofluorescence assays (IFA) showing DMT1-3×HA-tagged parasites stained with α-HA (DMT1, green) and DAPI (nuclear, blue), as well as antibodies or dyes: (**a**) MitoTracker Red CMXRos (mitochondria, red), (**b**) α-PfACP (apicoplast), (**c**) α-Rab5B (early endosome), (**d**) α-BiP (ER), (**e**) α-PfEXP2 (parasitophorous vacuolar membrane), (**f**) α-PfCRT (digestive vacuole membrane), (**g**) α-PfPM2 (digestive vacuole), (**h**) Nile Red (lipid bodies), or (**i**) α-Rab11A (post-Golgi). IFA images are also shown as 3D volume reconstructions (right-most column). Scale bars: 1 μm for IFA (black) or 0.5 μm for 3D volume reconstructions (white). **j**, Mean ± SEM percent DMT1-3×HA colocalization between DMT1 and the aforementioned organelles. Colocalization *p*-values (Student’s *t*-test): \**p* < 0.05; \*\**p* < 0.01; \*\*\**p* < 0.001; \*\*\*\**p* < 0.0001. Numbers of individual parasites analyzed and mean ± SEM and *p*-values for two different colocalization metrics (volume and Manders’ coefficient) are described in **Table S11**.

## References

1. World Health Organization. World malaria report 2024. 283 (2024).

2. Dhorda, M. et al. Artemisinin-resistant malaria in Africa demands urgent action. Science 385, 252–254 (2024).

3. Ishengoma, D.S. et al. Urgent action is needed to confront artemisinin partial resistance in African malaria parasites. Nat Med 30, 1807–1808 (2024).

4. Okombo, J. & Fidock, D.A. Towards next-generation treatment options to combat *Plasmodium falciparum* malaria. Nat Rev Microbiol 23, 178–191 (2024).

5. D’Alessandro, U. et al. Treatment of uncomplicated and severe malaria during pregnancy. Lancet Infect Dis 18, e133–e146 (2018).

6. Achan, J. et al. Quinine, an old anti-malarial drug in a modern world: role in the treatment of malaria. Malar J 10, 144 (2011).

7. Saeheng, T. & Na-Bangchang, K. Clinical pharmacokinetics of quinine and its relationship with treatment outcomes in children, pregnant women, and elderly patients, with uncomplicated and complicated malaria: a systematic review. Malar J 21, 41 (2022).

8. Cowman, A.F., Galatis, D. & Thompson, J.K. Selection for mefloquine resistance in *Plasmodium falciparum* is linked to amplification of the *pfmdr1* gene and cross-resistance to halofantrine and quinine. Proc Natl Acad Sci USA 91, 1143–7 (1994).

9. Reed, M.B., Saliba, K.J., Caruana, S.R., Kirk, K. & Cowman, A.F. Pgh1 modulates sensitivity and resistance to multiple antimalarials in *Plasmodium falciparum*. Nature 403, 906–909 (2000).

10. Sidhu, A.B., Verdier-Pinard, D. & Fidock, D.A. Chloroquine resistance in *Plasmodium falciparum* malaria parasites conferred by *pfcrt* mutations. Science 298, 210–213 (2002).

11. Ferdig, M.T. et al. Dissecting the loci of low-level quinine resistance in malaria parasites. Mol Microbiol 52, 985–97 (2004).

12. Nkrumah, L.J. et al. Probing the multifactorial basis of *Plasmodium falciparum* quinine resistance: evidence for a strain-specific contribution of the sodium-proton exchanger PfNHE. Mol Biochem Parasitol 165, 122–31 (2009).

13. Sanchez, C.P. et al. A HECT ubiquitin-protein ligase as a novel candidate gene for altered quinine and quinidine responses in *Plasmodium falciparum*. PLoS Genet 10, e1004382 (2014).

14. Dziekan, J.M. et al. Identifying purine nucleoside phosphorylase as the target of quinine using cellular thermal shift assay. Sci Transl Med 11, eaau3174 (2019).

15. Tinto, H. et al. Relationship between the *pfcrt* T76 and the *pfmdr-1* Y86 mutations in *Plasmodium falciparum* and *in vitro*/*in vivo* chloroquine resistance in Burkina Faso, West Africa. Infect Genet Evol 3, 287–92 (2003).

16. Sa, J.M. et al. Geographic patterns of *Plasmodium falciparum* drug resistance distinguished by differential responses to amodiaquine and chloroquine. Proc Natl Acad Sci USA 106, 18883–18889 (2009).

17. Patel, J.J. et al. Chloroquine susceptibility and reversibility in a *Plasmodium falciparum* genetic cross. Mol Microbiol 78, 770–87 (2010).

18. Veiga, M.I. et al. Globally prevalent PfMDR1 mutations modulate *Plasmodium falciparum* susceptibility to artemisinin-based combination therapies. Nat Commun 7, 11553 (2016).

19. Wicht, K.J., Mok, S. & Fidock, D.A. Molecular mechanisms of drug resistance in *Plasmodium falciparum* malaria. Annu Rev Microbiol 74, 431–454 (2020).

20. Takala-Harrison, S. & Laufer, M.K. Antimalarial drug resistance in Africa: key lessons for the future. Ann NY Acad Sci 1342, 62–7 (2015).

21. Ehrlich, H.Y., Jones, J. & Parikh, S. Molecular surveillance of antimalarial partner drug resistance in sub-Saharan Africa: a spatial-temporal evidence mapping study. Lancet Microbe 1, e209–e217 (2020).

22. Asua, V. et al. Changing pevalence of potential mediators of aminoquinoline, antifolate, and artemisinin resistance across Uganda. J Infect Dis 223, 985–994 (2021).

23. Wang, P., Read, M., Sims, P.F. & Hyde, J.E. Sulfadoxine resistance in the human malaria parasite *Plasmodium falciparum* is determined by mutations in dihydropteroate synthetase and an additional factor associated with folate utilization. Mol Microbiol 23, 979–86 (1997).

24. Cowman, A.F., Morry, M.J., Biggs, B.A., Cross, G.A. & Foote, S.J. Amino acid changes linked to pyrimethamine resistance in the dihydrofolate reductase-thymidylate synthase gene of *Plasmodium falciparum*. Proc Natl Acad Sci USA 85, 9109–13 (1988).

25. Peterson, D.S., Walliker, D. & Wellems, T.E. Evidence that a point mutation in dihydrofolate reductase-thymidylate synthase confers resistance to pyrimethamine in *falciparum* malaria. Proc Natl Acad Sci USA 85, 9114–8 (1988).

26. Fidock, D.A. et al. Mutations in the *P. falciparum* digestive vacuole transmembrane protein PfCRT and evidence for their role in chloroquine resistance. Mol Cell 6, 861–871 (2000).

27. Brenneman, K.V. et al. Optimizing bulk segregant analysis of drug resistance using *Plasmodium falciparum* genetic crosses conducted in humanized mice. iScience 25, 104095 (2022).

28. Mok, S. et al. Mapping the genomic landscape of multidrug resistance in *Plasmodium falciparum* and its impact on parasite fitness. Sci Adv 9, eadi2364 (2023).

29. Button-Simons, K.A. et al. The power and promise of genetic mapping from *Plasmodium falciparum* crosses utilizing human liver-chimeric mice. Commun Biol 4, 734 (2021).

30. Amaratunga, C. et al. Artemisinin-resistant *Plasmodium falciparum* in Pursat province, western Cambodia: a parasite clearance rate study. Lancet Infect Dis 12, 851–8 (2012).

31. Straimer, J. et al. K13-propeller mutations confer artemisinin resistance in *Plasmodium falciparum* clinical isolates. Science 347, 428–431 (2015).

32. Sidhu, A.B. et al. Decreasing *pfmdr1* copy number in *Plasmodium falciparum* malaria heightens susceptibility to mefloquine, lumefantrine, halofantrine, quinine, and artemisinin. J Infect Dis 194, 528–35 (2006).

33. Tripathi, A.K., Mlambo, G., Kanatani, S., Sinnis, P. & Dimopoulos, G. *Plasmodium falciparum* gametocyte culture and mosquito infection through artificial membrane feeding. J Vis Exp 161, 10.3791/61426 (2020).

34. Jiang, H. et al. High recombination rates and hotspots in a *Plasmodium falciparum* genetic cross. Genome Biol 12, R33 (2011).

35. Miles, A. et al. Indels, structural variation, and recombination drive genomic diversity in *Plasmodium falciparum*. Genome Res 26, 1288–1299 (2016).

36. Ranford-Cartwright, L.C. & Mwangi, J.M. Analysis of malaria parasite phenotypes using experimental genetic crosses of *Plasmodium falciparum*. Int J Parasitol 42, 529–34 (2012).

37. Su, X.Z. et al. A genetic map and recombination parameters of the human malaria parasite *Plasmodium falciparum*. Science 286, 1351–1353 (1999).

38. Amambua-Ngwa, A. et al. Chloroquine resistance evolution in *Plasmodium falciparum* is mediated by the putative amino acid transporter AAT1. Nat Microbiol 8, 1213–1226 (2023).

39. Amberg-Johnson, K. et al. Small molecule inhibition of apicomplexan FtsH1 disrupts plastid biogenesis in human pathogens. Elife 6, e29865 (2017).

40. Cowell, A.N. et al. Mapping the malaria parasite druggable genome by using *in vitro* evolution and chemogenomics. Science 359, 191–199 (2018).

41. Goodman, C.D., Uddin, T., Spillman, N.J. & McFadden, G.I. A single point mutation in the *Plasmodium falciparum* FtsH1 metalloprotease confers actinonin resistance. Elife 9, e58629 (2020).

42. Zhang, M. et al. Uncovering the essential genes of the human malaria parasite *Plasmodium falciparum* by saturation mutagenesis. Science 360, eaap7847 (2018).

43. The Pf3K Project: pilot data release 3. (2015).

44. Krogh, A., Larsson, B., von Heijne, G. & Sonnhammer, E.L. Predicting transmembrane protein topology with a hidden Markov model: application to complete genomes. J Mol Biol 305, 567–80 (2001).

45. Hoffman, K. & Stoffel, W. TMbase-a database of membrane spanning proteins segments. Biochim Biophys Acta 815, 468–476 (1993).

46. Jumper, J. et al. Highly accurate protein structure prediction with AlphaFold. Nature 596, 583–589 (2021).

47. Kim, J. et al. Structure and drug resistance of the *Plasmodium falciparum* transporter PfCRT. Nature 576, 315–320 (2019).

48. Liu, W. et al. Origin of the human malaria parasite *Plasmodium falciparum* in gorillas. Nature 467, 420–5 (2010).

49. Dziekan, J.M. et al. Cellular thermal shift assay for the identification of drug-target interactions in the *Plasmodium falciparum* proteome. Nat Protoc 15, 1881–1921 (2020).

50. Foley, M. & Tilley, L. Quinoline antimalarials: mechanisms of action and resistance and prospects for new agents. Pharmacol Ther 79, 55–87 (1998).

51. Straimer, J. et al. Site-specific genome editing in *Plasmodium falciparum* using engineered zinc-finger nucleases. Nat Methods 9, 993–8 (2012).

52. Palacpac, N.M. et al. Developmental-stage-specific triacylglycerol biosynthesis, degradation and trafficking as lipid bodies in *Plasmodium falciparum*-infected erythrocytes. J Cell Sci 117, 1469–80 (2004).

53. Jackson, K.E. et al. Food vacuole-associated lipid bodies and heterogeneous lipid environments in the malaria parasite, *Plasmodium falciparum*. Mol Microbiol 54, 109–22 (2004).

54. Hartwig, C.L. et al. Accumulation of artemisinin trioxane derivatives within neutral lipids of *Plasmodium falciparum* malaria parasites is endoperoxide-dependent. Biochem Pharmacol 77, 322–36 (2009).

55. Birnbaum, J. et al. A Kelch13-defined endocytosis pathway mediates artemisinin resistance in malaria parasites. Science 367, 51–59 (2020).

56. Gnadig, N.F. et al. Insights into the intracellular localization, protein associations and artemisinin resistance properties of *Plasmodium falciparum* K13. PLoS Pathog 16, e1008482 (2020).

57. Yang, T. et al. Decreased K13 abundance reduces hemoglobin catabolism and proteotoxic stress, underpinning artemisinin resistance. Cell Rep 29, 2917–2928 e5 (2019).

58. Ecker, A., Lehane, A.M., Clain, J. & Fidock, D.A. PfCRT and its role in antimalarial drug resistance. Trends Parasitol 28, 504–14 (2012).

59. Carter, M.D., Phelan, V.V., Sandlin, R.D., Bachmann, B.O. & Wright, D.W. Lipophilic mediated assays for beta-hematin inhibitors. Comb Chem High Throughput Screen 13, 285–92 (2010).

60. Sandlin, R.D. et al. Use of the NP-40 detergent-mediated assay in discovery of inhibitors of beta-hematin crystallization. Antimicrob Agents Chemother 55, 3363–9 (2011).

61. Olafson, K.N., Nguyen, T.Q., Rimer, J.D. & Vekilov, P.G. Antimalarials inhibit hematin crystallization by unique drug-surface site interactions. Proc Natl Acad Sci USA 114, 7531–7536 (2017).

62. Combrinck, J.M. et al. Insights into the role of heme in the mechanism of action of antimalarials. ACS Chem Biol 8, 133–7 (2013).

63. Bendrat, K., Berger, B.J. & Cerami, A. Haem polymerization in malaria. Nature 378, 138–9 (1995).

64. Wellems, T.E., Walker-Jonah, A. & Panton, L.J. Genetic mapping of the chloroquine-resistance locus on *Plasmodium falciparum* chromosome 7. Proc Natl Acad Sci USA 88, 3382–6 (1991).

65. Waller, K.L. et al. Chloroquine resistance modulated *in vitro* by expression levels of the *Plasmodium falciparum* chloroquine resistance transporter. J Biol Chem 278, 33593–601 (2003).

66. Sanchez, C.P. et al. The knock-down of the chloroquine resistance transporter PfCRT is linked to oligopeptide handling in *Plasmodium falciparum*. Microbiol Spectr 10, e0110122 (2022).

67. Tanveer, A. et al. An FtsH protease is recruited to the mitochondrion of *Plasmodium falciparum*. PLoS One 8, e74408 (2013).

68. Dahl, E.L. & Rosenthal, P.J. Multiple antibiotics exert delayed effects against the *Plasmodium falciparum* apicoplast. Antimicrob Agents Chemother 51, 3485–90 (2007).

69. Lewis, I.A. et al. Metabolic QTL analysis links chloroquine resistance in *Plasmodium falciparum* to impaired hemoglobin catabolism. PLoS Genet 10, e1004085 (2014).

70. Shafik, S.H. et al. The natural function of the malaria parasite’s chloroquine resistance transporter. Nat Commun 11, 3922 (2020).

71. Okombo, J. et al. Piperaquine-resistant PfCRT mutations differentially impact drug transport, hemoglobin catabolism and parasite physiology in *Plasmodium falciparum* asexual blood stages. PLoS Pathog 18, e1010926 (2022).

72. Howe, R., Kelly, M., Jimah, J., Hodge, D. & Odom, A.R. Isoprenoid biosynthesis inhibition disrupts Rab5 localization and food vacuolar integrity in *Plasmodium falciparum*. Eukaryot Cell 12, 215–23 (2013).

73. Kennedy, K. et al. Delayed death in the malaria parasite *Plasmodium falciparum* is caused by disruption of prenylation-dependent intracellular trafficking. PLoS Biol 17, e3000376 (2019).

74. Vaughan, A.M., et al. *Plasmodium falciparum* genetic crosses in a humanized mouse model. Nat Methods 12, 631–3 (2015).

75. Azuma, H. et al. Robust expansion of human hepatocytes in *Fah^-/-^/Rag2^-/-^/Il2rg^-/-^* mice. Nat Biotechnol 25, 903–10 (2007).

76. Li, H. & Durbin, R. Fast and accurate short read alignment with Burrows-Wheeler transform. Bioinformatics 25, 1754–60 (2009).

77. Cingolani, P. et al. A program for annotating and predicting the effects of single nucleotide polymorphisms, SnpEff: SNPs in the genome of *Drosophila melanogaster* strain *w^1118^*; *iso-2*; *iso-3*. Fly 6, 80–92 (2012).

78. Kanai, M. et al. Comparative analysis of *Plasmodium falciparum* genotyping via SNP detection, microsatellite profiling, and whole-genome sequencing. Antimicrob Agents Chemother 66, e0116321 (2022).

79. Li, H. et al. The sequence alignment/map format and SAMtools. Bioinformatics 25, 2078–2079 (2009).

80. de Hoon, M.J., Imoto, S., Nolan, J. & Miyano, S. Open source clustering software. Bioinformatics 20, 1453–1454 (2004).

81. Saldanha, A.J. Java Treeview--extensible visualization of microarray data. Bioinformatics 20, 3246–3248 (2004).

82. Mansfeld, B.N. & Grumet, R. QTLseqr: an R package for bulk segregant analysis with next-generation sequencing. Plant Genome 11 (2018).

83. Takagi, H. et al. QTL-seq: rapid mapping of quantitative trait loci in rice by whole genome resequencing of DNA from two bulked populations. Plant J 74, 174–83 (2013).

84. Magwene, P.M., Willis, J.H. & Kelly, J.K. The statistics of bulk segregant analysis using next generation sequencing. PLoS Comput Biol 7, e1002255 (2011).

85. Broman, K.W., Wu, H., Sen, S. & Churchill, G.A. R/qtl: QTL mapping in experimental crosses. Bioinformatics 19, 889–90 (2003).

86. Adjalley, S. & Lee, M.C.S. CRISPR/Cas9 editing of the *Plasmodium falciparum* genome. Methods Mol Biol 2470, 221–239 (2022).

87. Labun, K. et al. CHOPCHOP v3: expanding the CRISPR web toolbox beyond genome editing. Nucleic Acids Res 47, W171–W174 (2019).

88. Roy, K.R. et al. Multiplexed precision genome editing with trackable genomic barcodes in yeast. Nat Biotechnol 36, 512–520 (2018).

89. Fidock, D.A., Nomura, T. & Wellems, T.E. Cycloguanil and its parent compound proguanil demonstrate distinct activities against *Plasmodium falciparum* malaria parasites transformed with human dihydrofolate reductase. Mol Pharmacol 54, 1140–7 (1998).

90. Adjalley, S.H., Lee, M.C. & Fidock, D.A. A method for rapid genetic integration into *Plasmodium falciparum* utilizing mycobacteriophage Bxb1 integrase. Methods Mol Biol 634, 87–100 (2010).

91. Fidock, D.A. & Wellems, T.E. Transformation with human *dihydrofolate reductase* renders malaria parasites insensitive to WR99210 but does not affect the intrinsic activity of proguanil. Proc Natl Acad Sci USA 94, 10931–6 (1997).

92. Murithi, J.M. et al. The *Plasmodium falciparum* ABC transporter ABCI3 confers parasite strain-dependent pleiotropic antimalarial drug resistance. Cell Chem Biol 29, 824–839 e6 (2022).

93. Schindelin, J., et al. Fiji: an open-source platform for biological-image analysis. Nat Methods 9, 676–82 (2012).

94. Amos, B. et al. VEuPathDB: the eukaryotic pathogen, vector and host bioinformatics resource center. Nucleic Acids Res 50, D898–D911 (2022).

95. Sievers, F. et al. Fast, scalable generation of high-quality protein multiple sequence alignments using Clustal Omega. Mol Syst Biol 7, 539 (2011).

96. Needleman, S.B. & Wunsch, C.D. A general method applicable to the search for similarities in the amino acid sequence of two proteins. J Mol Biol 48, 443–53 (1970).

97. Madeira, F. et al. Search and sequence analysis tools services from EMBL-EBI in 2022. Nucleic Acids Res 50, W276–W279 (2022).

98. Goehring, A. et al. Screening and large-scale expression of membrane proteins in mammalian cells for structural studies. Nat Protoc 9, 2574–85 (2014).

99. Ncokazi, K.K. & Egan, T.J. A colorimetric high-throughput beta-hematin inhibition screening assay for use in the search for antimalarial compounds. Anal Biochem 338, 306–19 (2005).

100. Lopez-Barragan, M.J. et al. Directional gene expression and antisense transcripts in sexual and asexual stages of *Plasmodium falciparum*. BMC Genomics 12, 587 (2011).

101. Bushell, E. et al. Functional profiling of a *Plasmodium* genome reveals an abundance of essential genes. Cell 170, 260–272 (2017).

