## Supplementary_Figures for "*Plasmodium falciparum* quinine resistance is multifactorial and includes a role for the drug/metabolite transporters PfCRT and DMT1"

### Supplemental Figure 1

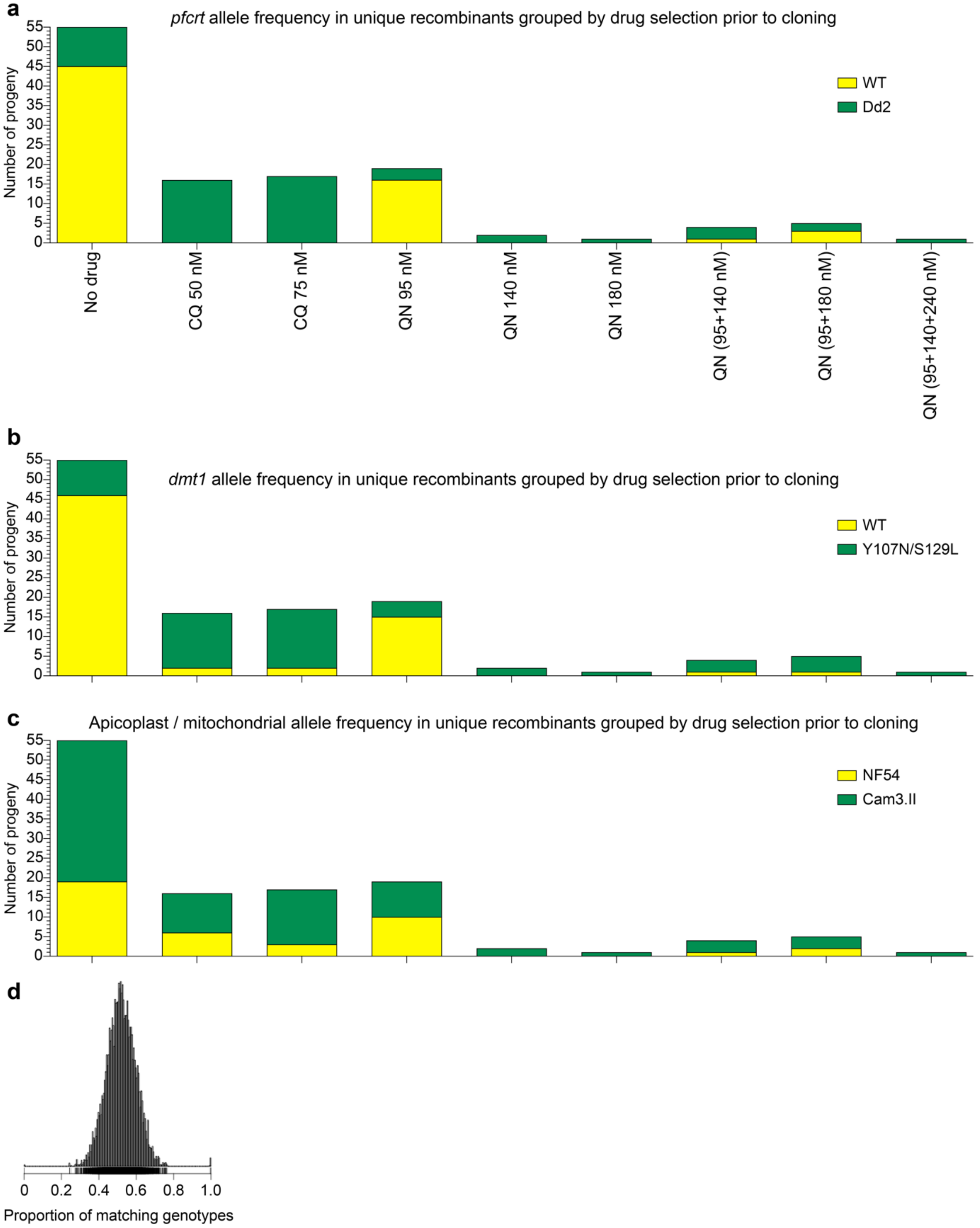

**Supplemental Fig. 1 | *pfcrt*, *dmt1*, and apicoplast/mitochondrial allele frequencies in the recombinant progeny.**

**a**-**c**, Number of unique recombinant progeny with the wild-type NF54 or mutant Cam3.II alleles for (**a**) *pfcrt,* (**b**) *dmt1*, and (**c**) the apicoplast and mitochondrial genomes. Data are grouped by the selection condition applied before obtaining the progeny clone. If a unique recombinant haplotype was obtained in more than one selection pressure condition, this was shown in every corresponding group. CQ, chloroquine; QN, quinine. The apicoplast and mitochondrial haplotypes are: Cam3.II: *PF3D7_API00100*, S25G; *PF3D7_API02100*, V46I; *PF3D7_API02300*, K118R; *PF3D7_API03600*, L195; *PF3D7_API04400*, S658; *mal_mito_1*, wild-type; NF54: wild-type for all apicoplast genes; *mal_mito_1*, I250V. **d**, Genetic relatedness between the 120 recombinant progeny clones and two parents was measured by identity-by-descent in multiple pairwise comparisons.

#### Supplemental Figure 2

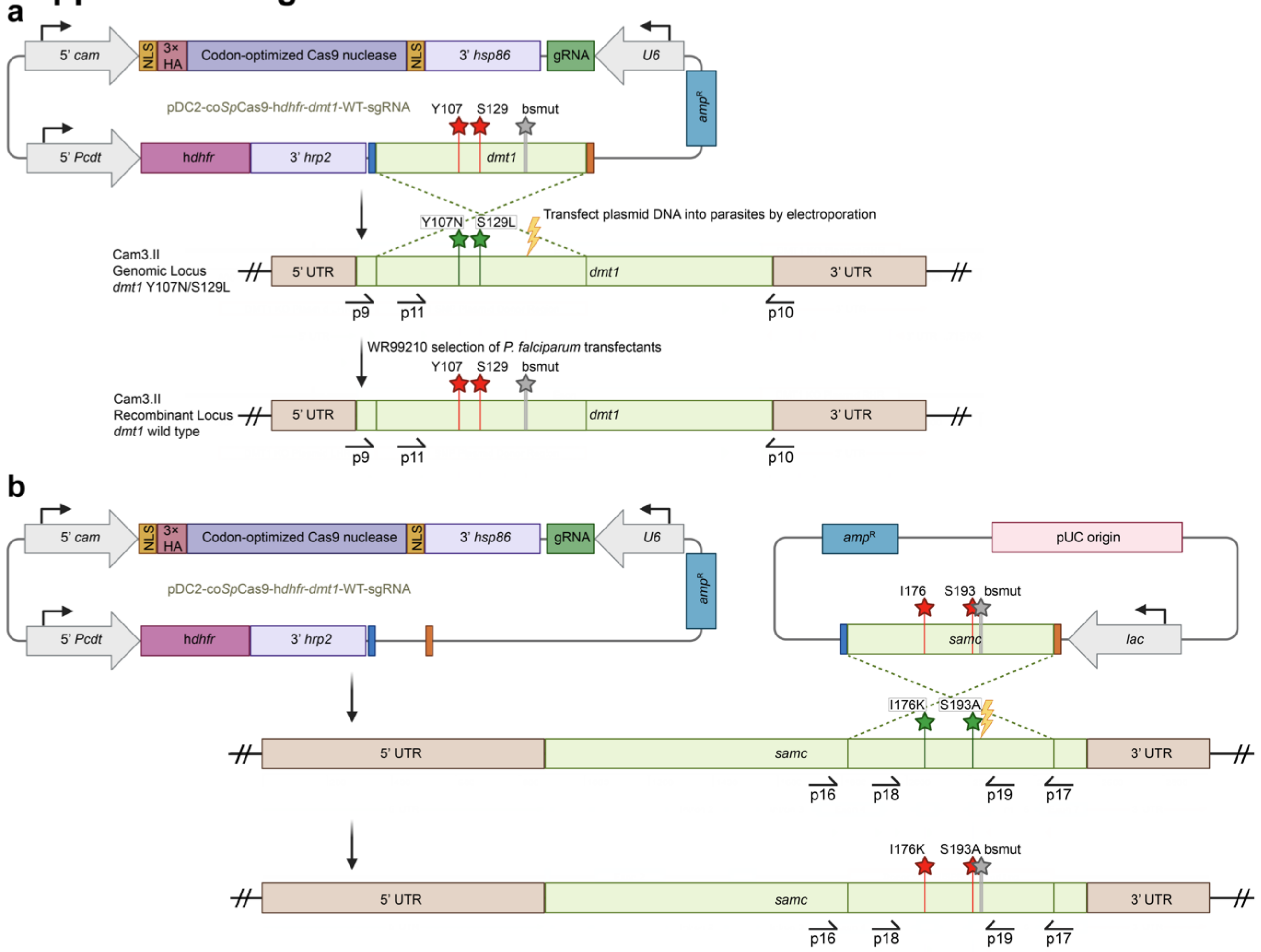


**Supplemental Fig. 2 | CRISPR/Cas9 plasmid construction and editing strategy for single nucleotide polymorphism (SNP) editing.**

**a**, All-in-one plasmid approaches used for CRISPR/Cas9-mediated SNP editing of QTL candidate genes in the cross parents and progeny, with the *dmt1* gene shown as a representative. This plasmid contained the *cam* promoter to express codon-optimized Cas9 and a *PcDT* promoter to express the WR99210-selectable h*dhfr* marker. This schematic shows a door fragment encoding the two *dmt1* wild-type codons Y107 and S129 along with binding-site mutations (bsmut; shown as grey stars) to prevent further cleavage of gene-edited parasites. Plasmid DNA was transfected into parasites by electroporation to initiate a double-stranded break, and subsequent homology-directed repair and replacement of the mutated codons encoding Y107N and S129L (green stars) with the wild-type sequence (red stars). Donor plasmids were also designed with mutant codons that were gene edited into parasites with wild-type *dmt1*. **b**, The two-plasmid strategy used for CRISPR/Cas9-mediated SNP editing of *samc*. The donor for SAMC was kept in the pUC-GW-*amp* vector and co-transfected with a Cas9 plasmid expressing the gRNA. *cam*, calmodulin; gRNA, guide RNA; h*dhfr*, human dihydrofolate reductase; *hrp2*: histidine-rich protein 2; *hsp86*, heat shock protein 86; *PcDT*, *P. chabaudi* dihydrofolate reductase-thymidylate synthase; *amp*$^{R}$, ampicillin resistance gene; *dmt1*, *P. falciparum* drug/metabolite transporter 1; UTR, untranslated region. Primers used for cloning and verification are described in **Table S8**. Plasmids are described in **Table S9**.

### Supplemental Figure 3

a

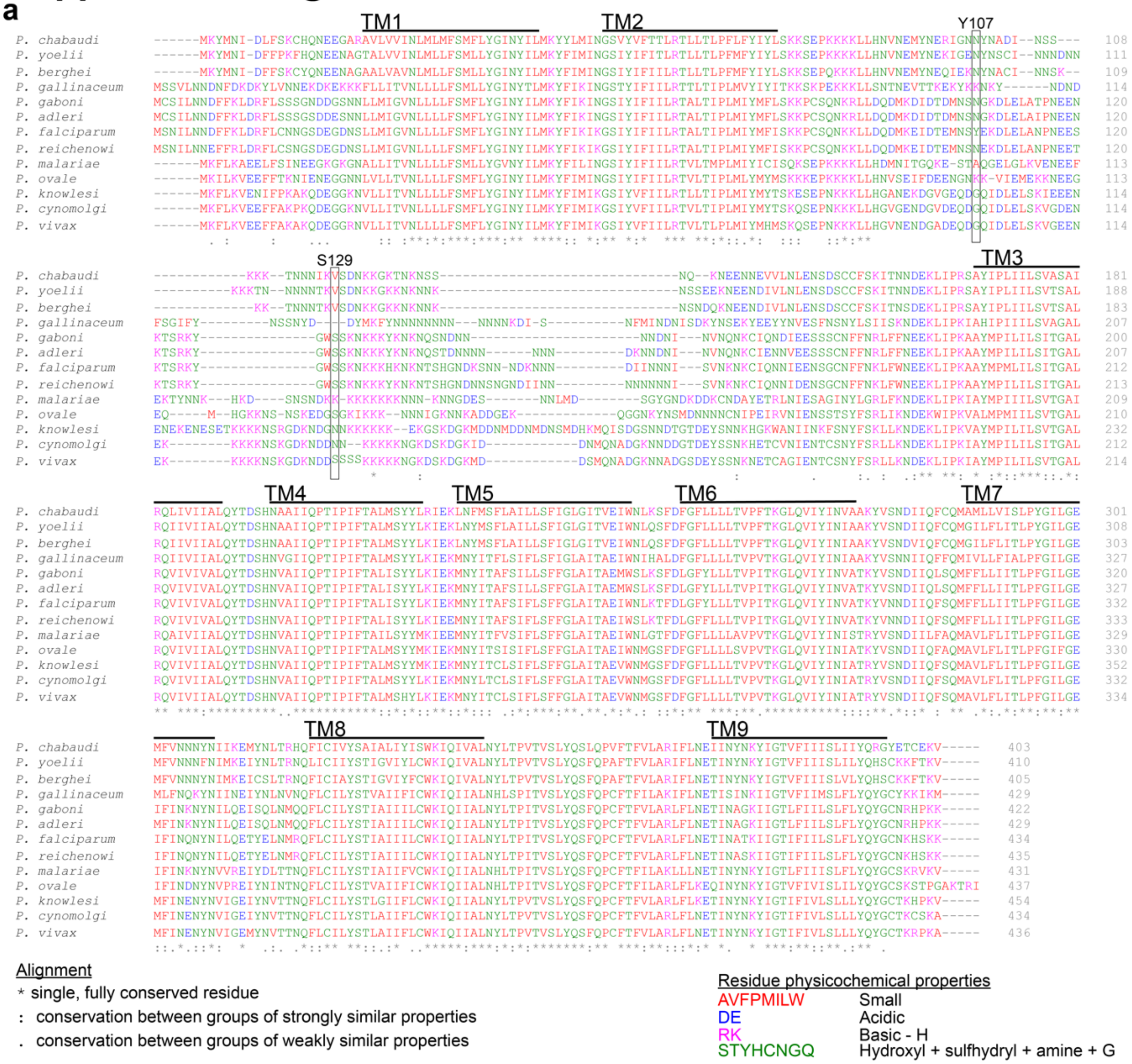


b

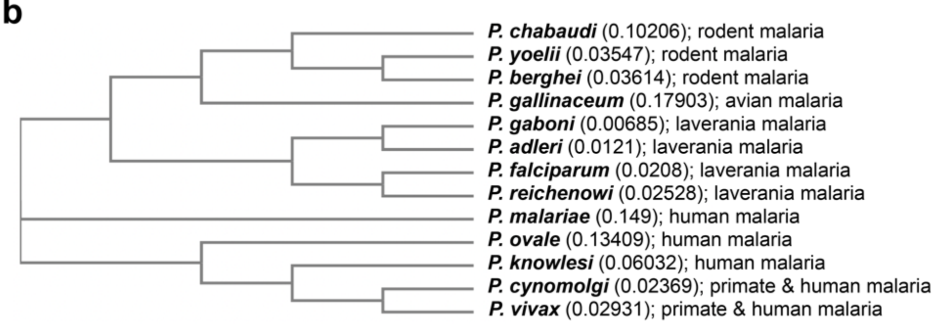

**Supplemental Fig. 3 | Residues in predicted transmembrane regions of PfDMT1 show conservation between orthologues of DMT1 in other *Plasmodium spp.***

**a**, Multiple sequence alignment of NF54 *P. falciparum* DMT1 with sequences of twelve orthologs of DMT1 sequences in rodent (*P. chabaudi*: chabaudi, *P. yoelii*: 17X, *P. berghei*: ANKA), avian (*P. gallinaceum*: 8A), laverania (*P. gaboni*: SY75, *P. adleri*: G01, *P. reichenowi*: CDC), human (*P. malariae*: UG01, *P. ovale*: GH01, *P. knowlesi*: H), and primate / human (*P. cynomolgi*: M, *P. vivax*: P01) reference *Plasmodium* species. The residues corresponding to the Y107N and S129L mutations in *P. falciparum* Cam3.II, and the TMHMM-predicted transmembrane domains of PfDMT1 (dark line) are indicated. **b**, Phylogenetic tree of the DMT1 sequences from the 13 *Plasmodium spp.* based on their multiple sequence alignment, with the tree "lengths" shown in parentheses (indicative of the evolutionary distance and genetic change between sequences).

#### Supplemental Figure 4

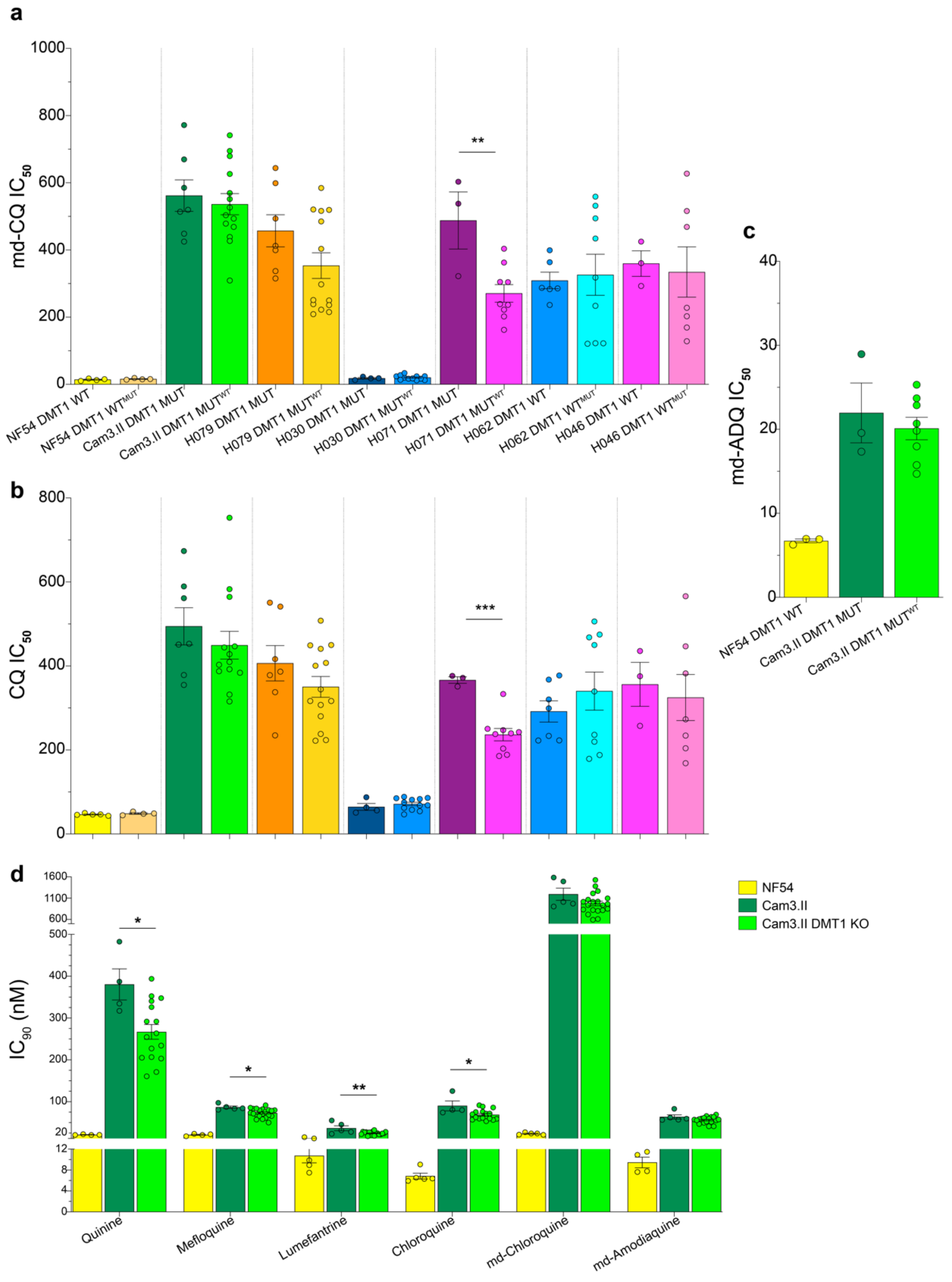

**Supplemental Fig. 4 | Phenotypic responses of DMT1 gene-edited parents and progeny to monodesethyl-chloroquine (md-CQ), chloroquine (CQ), and monodesethyl-amodiaquine (md-ADQ).**

**a**-**c**, md-CQ (**a**), CQ (**b**), and md-ADQ (**c**) mean ± SEM $IC_{50}$ values. For each parasite background, we phenotyped a parental unedited parasite (endogenous DMT1 haplotype) (dark color) and its isogenic gene-edited line (with binding-site mutations and Y107N / S129L mutant or a wild-type revertant) (light bright color). *p* values (Student's *t*-test) are indicated (N=3–14 with technical duplicates). **d**, Quinine, mefloquine, lumefantrine, CQ, md-CQ, and md-ADQ response in Cam3.II DMT1 knockout parasites as measured by mean ± SEM $IC_{90}$ values. *p*-values (Student's *t*-test) are indicated for the Cam3.II KO strain vs the Cam3.II parent (N=4–21, with technical duplicates). $^{*}p < 0.05$, $^{**}p < 0.01$, $^{***}p < 0.001$. WT: wild-type; MUT: Y107N / S129L mutation.

#### Supplemental Figure 5

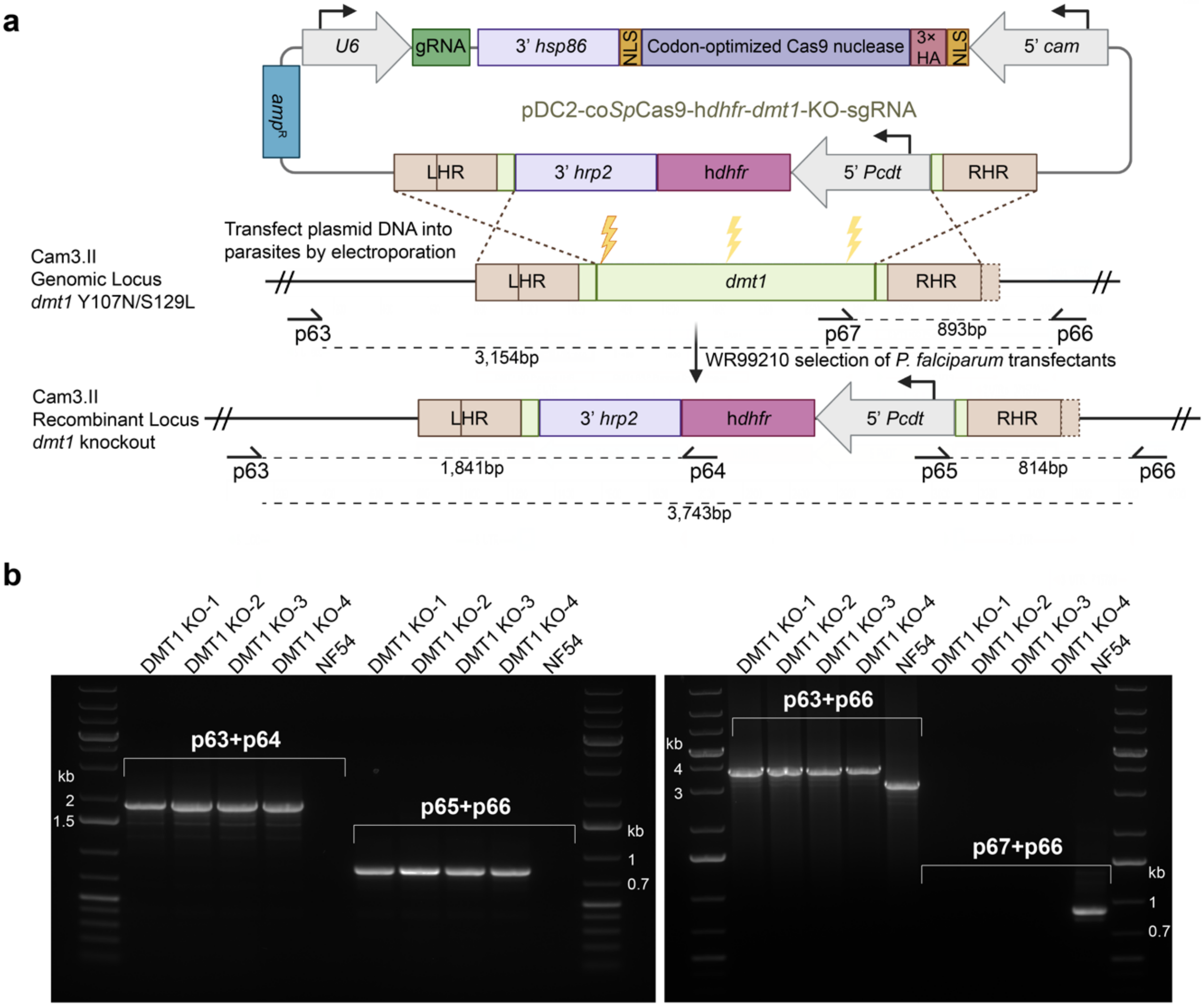


**Supplemental Fig. 5 | CRISPR/Cas9-based generation of DMT1 knockout Cam3.II parasites.** The all-in-one CRISPR/Cas9-based DMT1 knockout tagging strategy. **a**, CRISPR/Cas9 plasmids included a *dmt1* left homology region (LHR) and right homology region (RHR) flanking a human dhfr selectable marker cassette. Expression of Cas9 and a gRNA led to a double-stranded break in *dmt1*, which was repaired using the LHR and RHR homologous regions. This resulted in introduction of the selectable marker cassette (selected using 2.5 nM WR99210) and a dmt1 knockout (KO). Three plasmids were generated, each with its own gRNA (yellow thunderbolts) that targeted an internal *dmt1* sequence that was deleted during the gene editing event. **b**, PCR and gel electrophoresis-based verification of complete *dmt1* knockout in three Cam3.II parasite replicates, compared to a NF54 unedited control. Results confirmed the presence of the 5'-incorporated h*dhfr* cassette (p63+p64), 3'-incorporated *PcDT* promoter (p65+p66), ~600 bp larger size of the DMT1 recombinant knockout locus (p63+p66), and lack of the deleted *dmt1* locus (p67+p66). PCR primers and expected PCR product sizes are indicated above in (**a**). Primers used for verification and cloning are described in **Table S8**. Plasmids are described in **Table S9**.

#### Supplemental Figure 6

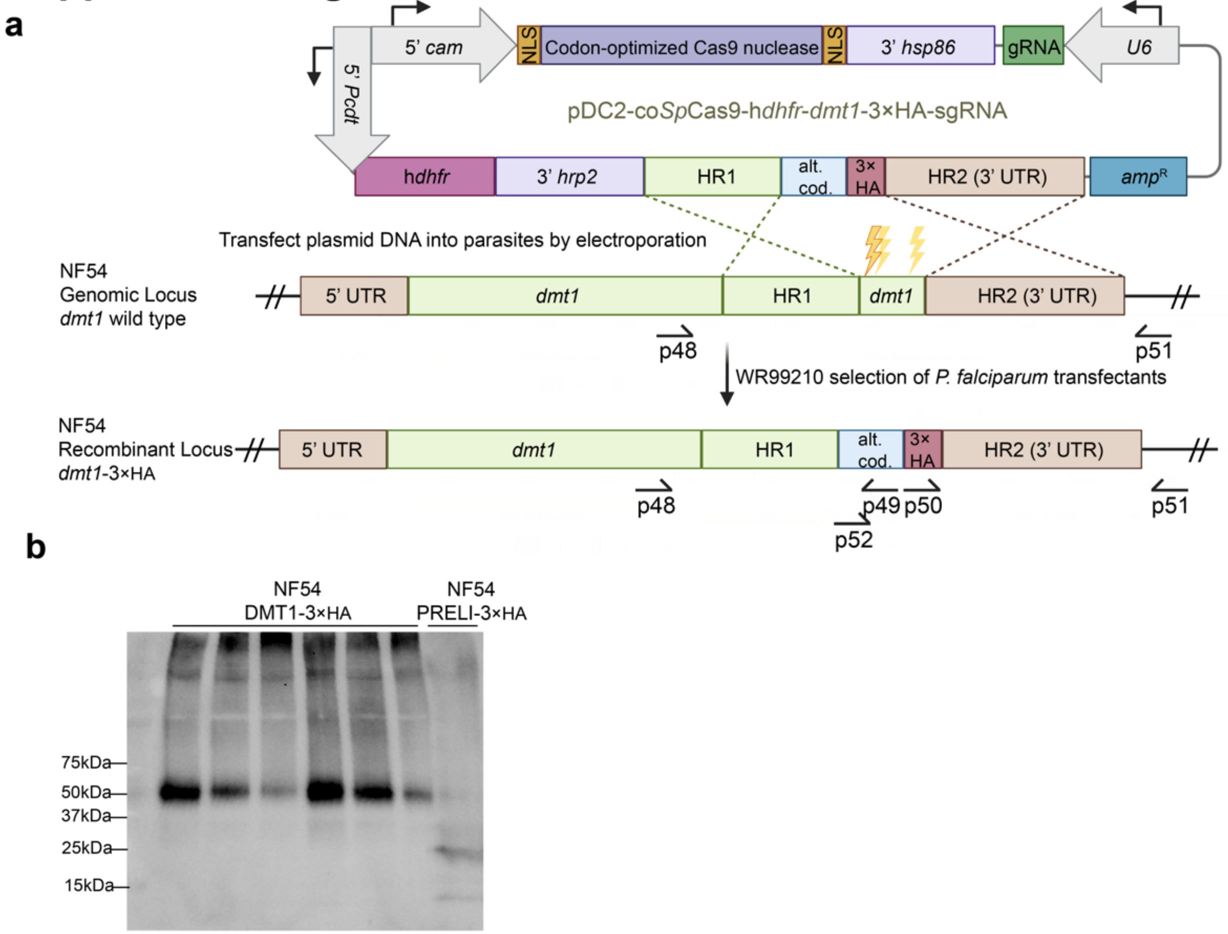


**Supplemental Fig. 6 | CRISPR/Cas9-based generation of DMT1 3' 3×HA-tagged NF54 parasites.**

**a**, The all-in-one CRISPR/Cas9-based DMT1 3' 3×HA tagging strategy. CRISPR/Cas9 plasmids were constructed as depicted in Supplemental Fig. 3, except the donor template included a homology region 1 (HR1) (342 bp), a recodonized 3' end of *dmt1* to disrupt homology with no stop codon (165 bp), a filler sequence (6 bp), a 3×HA epitope tag followed by a stop codon, and a 3' untranslated region homology region 2 (HR2; 509 bp). Three plasmids were generated, each with a separate gRNA located within the recodonized region. Thus, binding-site mutations were not necessary. **b**, Western blot of NF54 DMT1-3×HA (showing an expected 50 kDa band) and NF54 PRELI-3×HA parasites as a 3×HA positive control (~25 kDa band expected). Parasite lysates were incubated with an anti-HA antibody. Primers used for cloning and verification are described in **Table S8**. Plasmids are described in **Table S9**.
